# The 5’ fusion partner modulates response to the STAMP inhibitor asciminib in *ABL*-rearranged ALL

**DOI:** 10.1101/2025.05.06.652078

**Authors:** Laura N Eadie, Daniel P McDougal, Elias Lagonik, Caitlin E Schutz, Tace S Conlin, Elyse C Page, Jacqueline Rehn, John B Bruning, Andrew S Moore, David T Yeung, Timothy P Hughes, Deborah L. White

## Abstract

*ABL*-rearranged (*ABL*r) acute lymphoblastic leukaemia (ALL) is associated with high rates of treatment failure and relapse and novel treatments are required. We investigated activity of the STAMP inhibitor asciminib in non-*BCR::ABL1 ABL*r ALL. The most common fusion in *ABL*r ALL is *NUP214::ABL1*, which is associated with aggressive disease. For the first time we establish asciminib activity in three pre-clinical patient derived xenograft models of *NUP214::ABL1* ALL. Treatment with asciminib reduced *NUP214::ABL1* leukaemic burden, splenomegaly and ABL1 kinase activation. We observed significantly increased survival outcomes in asciminib-treated versus control mice. Additionally, site directed mutagenesis, *in vitro* cell death assays and *in silico* structural modeling defined a region of the ABL1 SH3 domain critical for asciminib efficacy and necessary for mediation of allosteric inhibition. Our findings establish asciminib as a potential treatment for *NUP214::ABL1* ALL, significantly expanding the number of ALL patients who may benefit from asciminib therapy which has an excellent safety and tolerability profile.

## Introduction

Acute lymphoblastic leukemia (ALL) is predominantly B-cell in origin, with T-cell ALL (T-ALL) representing 15% to 20% of ALL diagnoses depending on age group^1,2^. Survival rates for children with ALL now approach 90% in developed countries^3^, however, children with high-risk disease, neonates, adolescents and young adults and older adults still have poor outcomes and increased risk of relapse^4^. For those patients who do relapse, long term survival rates are dismal, particularly in the T-ALL setting^5^. A large subset of B-cell ALL (B-ALL) patients have a gene expression signature similar to that of *BCR::ABL1*+ (Ph+) leukemia, harboring gene fusions involving kinase and cytokine receptor genes^6^. This subtype is commonly known as Philadelphia (Ph)-like ALL, which is now recognized as a high-risk disease entity in the 2022 World Health Organisation’s classification of acute leukemias^7^.

Up to 20% of Ph-like ALL encompasses *ABL*-class fusions which frequently involve fusion of the *ABL1* gene to various partners including *NUP214*, *ETV6*, *RANBP2, RCSD1, SFPQ*, *ZMIZ1, SNX2* and others (collectively ∼30%)^8–12^. The *NUP214::ABL1* fusion is also reported in up to 6% of T-ALL cases depending on the age group^13–16^. Patients with *ABL*-rearranged (*ABL*r) disease experience very poor early response, event free (EFS) and overall survival (OS)^17^. The majority of *ABL1* gene fusions result from either an *ABL1* exon 2 (*ETV6*, *RANBP2*, *ZMIZ1*) or exon 4 (*RCSD1*, *SFPQ*, *SNX2*) breakpoint. However, the *NUP214::ABL1* fusion has been reported with multiple *ABL1* breakpoints. The *ABL1* exon 2 breakpoint has been identified in several T-ALL cohorts^15,18–20^ while *ABL1* exon 3 breakpoints have been identified in B-ALL patients^11,21,22^ and one T-ALL patient^16^. Recently, rare breakpoints in *ABL1* exons 1 and 4 have been identified in 3/27 *NUP214::ABL1* cases from a cohort of >1300 T-ALL patients^16^. The *NUP214::ABL1* gene fusion is a leukemic driver resulting in aggressive disease with poor outcomes in both the B- and T-ALL settings^16–18,23^.

Early *in vitro* and *in vivo* studies demonstrated that ABL ATP-competitive tyrosine kinase inhibitors (TKIs) such as dasatinib, used as front-line treatment in chronic myeloid leukemia (CML), are effective against Ph-like fusions involving *ABL1* or *ABL2*^11,24–28^. The TKIs imatinib or dasatinib are not routinely used in the treatment of non-*BCR::ABL1 ABL*r patients, however, recent studies suggest *ABL*r patients receiving concomitant TKI therapy during consolidation chemotherapy have better early responses potentially leading to better long-term outcomes^17,29^. Indeed, clinical trials evaluating the efficacy of TKIs in combination with chemotherapy in patients with Ph+ and ABL-class Ph-like ALL (NCT06124157, ISRCTN #64515327) demonstrated improved long-term outcomes compared with patients not receiving TKI^29,30^. In addition, inclusion of TKIs in the treatment regimen for ALL patients with the *BCR::ABL1* fusion, has reduced the need for hematopoietic stem cell transplantation (HSCT) in a proportion of this cohort and the associated high treatment related mortality^31–33^. However, there are cases where development of ATP-pocket point mutations, that prevent effective TKI binding, have led to TKI-resistance following an initial successful response^34–37^. Additionally, dasatinib and ponatinib are associated with significant side effects and off target interactions^38^ and new treatment strategies are required to overcome TKI failure and avoid potential long-term effects.

To overcome resistance to ATP-competitive TKIs^39^, a novel targetable binding pocket within the ABL kinase was identified^40,41^. The myristate binding pocket located in the C-lobe of the ABL kinase domain, is an allosteric negative regulator of ABL^42^ when bound by the myristate moiety present on native ABL. However, fusion of *ABL1* to *BCR* or other 5’ partner genes truncates the myristate moiety preventing autoinhibition and resulting in the constitutively active kinase and downstream signaling observed in *ABL*r leukemia patients^42^.

Asciminib is a first-in-class Specifically Targeting the ABL Myristoyl Pocket (STAMP) compound that mimics the myristate moiety, binding within the pocket with high affinity. Following binding of asciminib, the SH3 and SH2 domains bind to and stabilize the kinase domain, locking ABL1 in the inactive conformation^43^. Asciminib has recently been approved as a front-line therapy for newly diagnosed adults with CML as well as those who have previously been treated with TKIs^39,44^. A trial assessing TKI-asciminib combination therapy in Ph+ ALL patients has recently concluded (NCT02081378) with early results demonstrating durable molecular responses^45^. Importantly, asciminib has demonstrated superior efficacy in *de novo* CML patients compared with TKIs and has an excellent safety and tolerability profile^44^.

Initial *in vitro* studies examining potential mechanisms of asciminib resistance reported the importance of the ABCG2 efflux transporter and myristoyl pocket point mutations^46,47^. More recent investigations focused on the allosteric action of asciminib and the role of the ABL SH3 and SH2 domains for asciminib activity in *in vitro* models of CML^48,49^. We recently reported the efficacy of asciminib in *ABL*r ALL cell lines and highlighted that asciminib efficacy was dependent on inclusion of a critical region of *ABL* SH3 domain^50,51^. Others subsequently demonstrated that the exon 2-encoded part of the SH3 domain is essential for asciminib efficacy against *BCR::ABL1* in the CML setting^48,49^.

In this current study, we comprehensively evaluate asciminib efficacy in three pre-clinical patient derived xenograft (PDX) models of *ABL*r B- and T-ALL reporting sensitivity of *NUP214::ABL1* disease for the first time. We also use cell lines engineered to express different regions of the ABL1 SH3 and SH2 domains to investigate the key region of ABL1 required for asciminib efficacy using *NUP214::ABL1* as the model gene fusion. Results demonstrate varying asciminib sensitivity in cell death assays following deletion of specific regions of the SH3 domain. We then use AlphaFold3^52^ structural modeling to investigate the functional outcomes when different regions of the SH3 and SH2 domains are deleted. Our findings help to better characterize ABLr ALL patient populations who may benefit from the addition of asciminib to treatment regimens.

## Results

### Asciminib treatment significantly increases survival outcomes in PDX models of *ABL*r ALL

Following our recent discovery of asciminib activity in models of *NUP214::ABL1* ALL *in vitro*^50^, we investigated asciminib efficacy in PDX models of disease. We established three PDX models from cells of one T-ALL patient and two B-ALL patients with a *NUP214::ABL1* fusion (Figure S1; Table S1). Leukemic engraftment was confirmed by human (h) CD45+ cells in murine peripheral blood (PB). Engraftment levels were also confirmed by hCD7+ and hCD19+ cells for T-ALL and B-ALL PDX, respectively (data not shown). Once a threshold level of 5% hCD45+ cells was observed in the PB, treatment was commenced. Vehicle control mice developed an aggressive leukemia; conversely, asciminib treatment significantly delayed disease expansion in all PDX models (Figure 1A-B; Figure S2).

**Figure 1.**
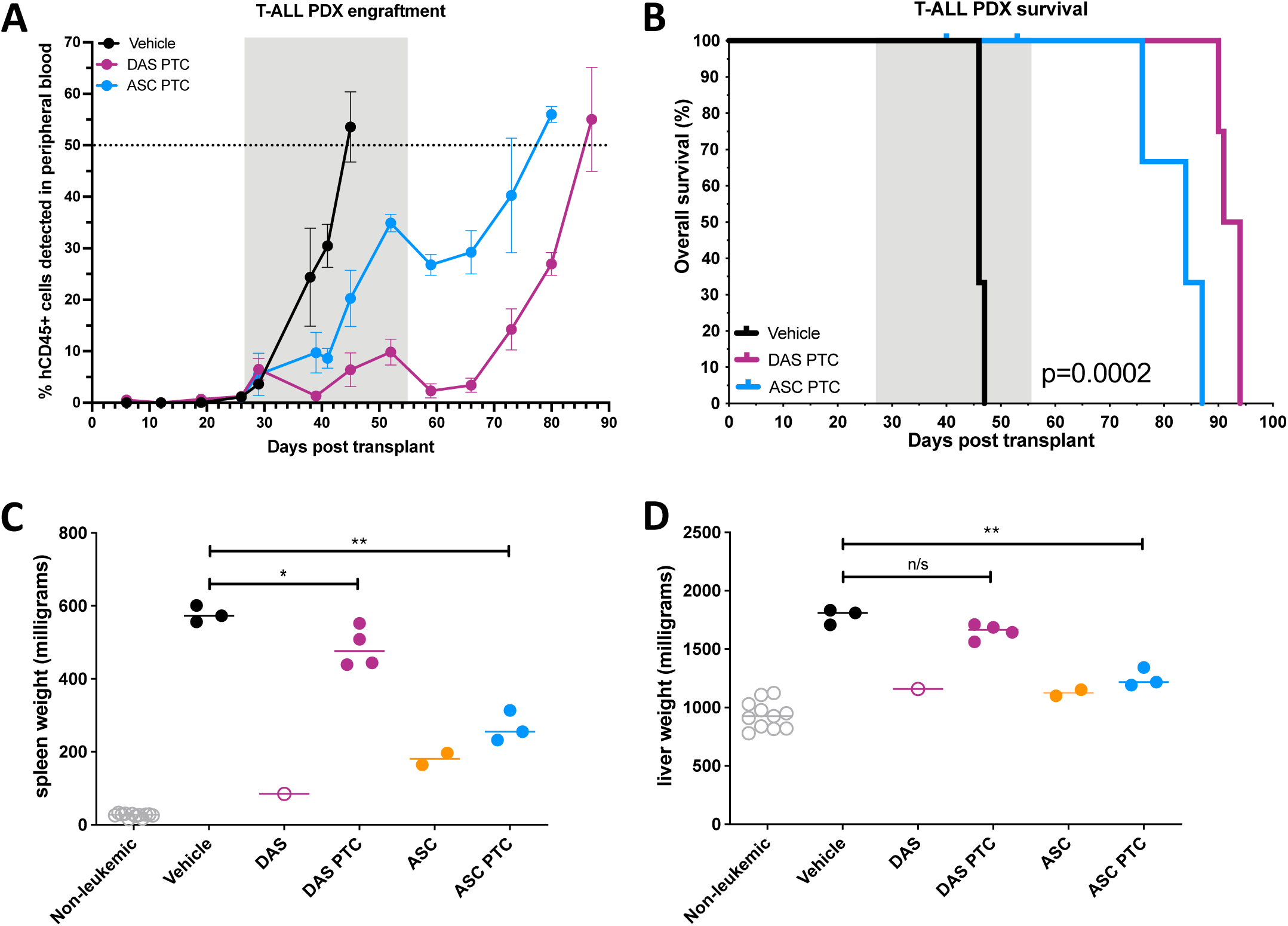
Treatment with asciminib show similar efficacy to dasatinib in a T-ALL *NUP214::ABL1* mouse model. A patient derived xenograft (PDX) was established from a T-ALL patient with *NUP214::ABL1* disease. A) Engraftment of leukemic cells was tracked via measurement of hCD45+ cells in the peripheral blood. Points on the graphs represent mean ±SD. Treatment windows are highlighted in grey. B) Kaplan-Meier curves determined survival with censored mice culled prior to end of treatment window shown as vertical lines. Censored mice were excluded from survival analyses. Significance was evaluated between all curves and determined by Log-rank test. C) Spleen or D) liver weights of mice at experimental endpoint, either on treatment or post-treatment cessation. Values represent mean ± SD, horizontal lines represent the median. Welch’s unpaired t-test calculated significance *p<0.05, **p<0.01. DAS=dasatinib, ASC=asciminib, PTC=post-treatment cessation.

To assess the effect of treatment on peripheral organ infiltration, mice were divided into two cohorts: mice in cohort 1 were humanely euthanized immediately following completion of inhibitor treatment and those in cohort 2 upon clinical signs of illness or when PB levels of hCD45+ reached 50%. Mice in cohort 2 were termed the post-treatment cessation (PTC) cohort and were assessed for alterations in survival outcomes. *NUP214::ABL1* T-ALL mice treated with asciminib demonstrated significantly improved survival compared with vehicle controls (experimental endpoint at day 84 vs 46; *p*=0.0100; Figure 1B). As expected, dasatinib treatment also significantly increased survival compared with vehicle (93 vs 46 d, *p*=0.0046). Importantly, survival for asciminib-treated mice was comparable to that of dasatinib-treated mice. Similar observations were made for both B-ALL PDX models (asciminib vs vehicle Patient 1=110 vs 74 d and Patient 2=103 vs 76 d; both *p*=0.0067; Figure S2B,D). Taken together, these results demonstrate the potential of asciminib as a therapy for *NUP214::ABL1* ALL patients.

Following asciminib treatment completion, we observed a decrease in peripheral organ weights for cohort 1 mice compared with vehicle mice. Spleen and liver weights of *NUP214::ABL1* T-ALL mice culled while receiving either dasatinib (n=1, 85 mg and 1158 mg, respectively) or asciminib (n=2, 181±23 mg and 1126±37 mg, respectively) treatment were equivalent to that of healthy mice (Figure 1C,D). Analogous observations were made for *NUP214::ABL1* B-ALL mice where asciminib treatment significantly reduced spleen (Patient 1 cohort, *p*=0.0390; Patient 2 cohort, *p*=0.0109) and liver (Patient 2 cohort, *p*=0.0357) weights compared with vehicle-treated mice (Figure S3A-D).

However, in all three PDX models, we observed an increase in organ weights following treatment cessation (Figure 1C,D; Figure S3A-D). Interestingly, in the *NUP214::ABL1* T-ALL model, even when PB levels of hCD45+ cells reached >50%, spleen and liver weights remained significantly lower in mice treated with asciminib compared with vehicle (267±42 vs 577±28 mg, *p*=0.0013 and 1250±80 vs 1784±67 mg, *p*=0.0011, respectively). This indicated a sustained effect of asciminib which was not observed in dasatinib-treated mice (Figure 1C,D).

### Asciminib treatment of *NUP214::ABL1* mice reduces leukemic blast density in bone marrow and peripheral organs

To examine the extent of leukemic infiltration, flow cytometry for hCD7+ (T-ALL PDX) or hCD19+ (B-ALL PDXs), RT-PCR specific for the *NUP214::ABL1* fusions and/or hematoxylin and eosin (H&E) or hCD45 histology were utilized (Figures 2,S4-8 and summarized in Table 1). Asciminib treatment reduced the level of circulating leukemic blasts in all PDX models compared with vehicle treated cohorts and normalized WBC counts (Supplementary Note 1). Flow cytometric analyses and H&E histology demonstrated mice culled while undergoing dasatinib or asciminib treatment had decreased levels of leukemic blasts in the bone marrow and peripheral organs compared with the vehicle-treated controls (Figure 2A, Figures S4,S6-8). Additionally, H&E histology of on-treatment liver sections demonstrated reduced blast island formation compared with the number of blast islands in control samples (Figure2B; Figures S7,S8). Additionally, H&E histology of on-treatment liver sections demonstrated reduced blast island formation compared with the number of blast islands in control samples (Figure2B; Figures S7,S8). While splenic blast density was not visibly decreased during asciminib treatment, there was a reduction in the overall blast number due to the significantly reduced spleen size of asciminib-treated mice (Figures S6-S8). The blast population density in the BM and peripheral organs increased following treatment cessation compared with respective on-treatment samples (Figure 2; Figures S6-S8).

**Figure 2.**
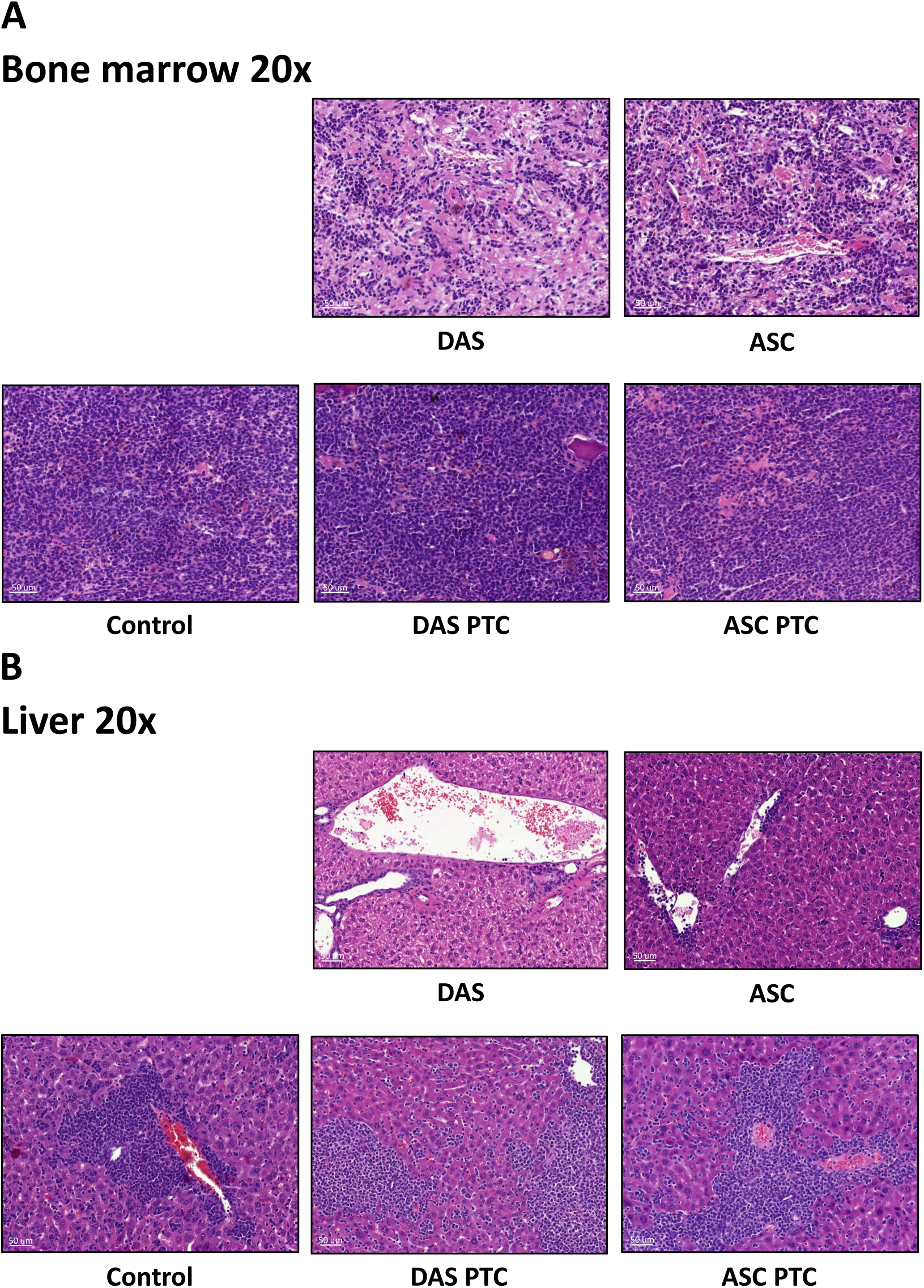
Characterization of leukemic infiltration in a *NUP214::ABL1* PDX model of T-ALL at experimental endpoint, either on treatment or post-treatment cessation. Organ sections stained with hematoxylin and eosin from *NUP214::ABL1* PDX mice. Images of **A)** bone marrow **B)** liver. Scale bars and magnifications are indicated. Representative images of sections from n=3 mice. DAS=dasatinib, ASC=asciminib, PTC=post-treatment cessation.

**Table 1.** Confirmation of disease in bone marrow and peripheral organs from T-ALL and B-ALL PDX models.

| **T-ALL Patient** | **BM** | **SPL** | **LIV** | **THY** |
| --- | --- | --- | --- | --- |
| **Vehicle control** | Y (FC,H) | Y (FC,PCR,H) | Y (FC,PCR,H) | Y (FC,H) |
| **DAS** | Y (FC<sup>lo</sup>,H) | Y (FC,PCR,H) | Y (FC,PCR,H) | Y (H) |
| **DAS PTC** | Y (FC,H) | Y (FC,PCR,H) | Y (FC,PCR,H) | Y (FC,PCR,H) |
| **ASC** | Y (FC<sup>lo</sup>,PCR,H) | Y (FC,PCR,H) | Y (FC,PCR,H) | Y (H) |
| **ASC PTC** | Y (FC,H) | Y (FC,PCR,H) | Y (FC,PCR,H) | Y (FC,PCR,H) |
| **B-ALL Patient 1** | **BM** | **SPL** | **LIV** | **THY** |
| **Vehicle control** | Y (FC,PCR) | Y (FC,PCR,H) | Y (FC,PCR,H) | Y (FC) |
| **ASC** | Y (FC,PCR) | Y (FC,PCR,H) | Y (FC,PCR,H) | NTE |
| **ASC PTC** | Y (FC,PCR) | Y (FC,PCR,H) | Y (FC,PCR,H) | Y (H) |
| **B-ALL Patient 2** | **BM** | **SPL** | **LIV** | **THY** |
| **Vehicle control** | Y (FC,PCR) | Y (FC,PCR,H) | Y (FC,PCR,H) | Y (H) |
| **ASC** | Y (FC,PCR) | Y (FC,PCR,H) | Y (FC,PCR,H) | NTE |
| **ASC PTC** | Y (FC,PCR) | Y (FC,PCR,H) | Y (FC,PCR,H) | Y (FC,PCR,H) |

hCD7/hCD19+ flow cytometry and/or *NUP214::ABL1* fusion RT-PCR and/or H&E or hCD45 histology was performed on a minimum of 3 samples. DAS=dasatinib, ASC=asciminib, PTC=post treatment cessation, BM=bone marrow, SPL=spleen, LIV=liver, THY=thymus, Y=yes, N=no, nd=not determined, NTE=no thymus evident, FC=flow cytometry, H=histology.

### Phosphorylation of STAT5 is reduced with asciminib treatment

Activation of STAT5 signalling, and asciminib-mediated inhibition, was demonstrated in our *in vitro* models^50^ and analogous observations were confirmed *in vivo* (Figure 3A). In the *NUP214::ABL1* T-ALL PDX, high levels of phosphorylated (p) STAT5 (MFI: 288±70) were detected in cells from vehicle-treated mice. Treatment with asciminib abrogated STAT5 pathway activation (MFI pSTAT5: 117±5). Once mice reached experimental endpoint post-treatment cessation, cells were harvested and STAT5 phosphorylation re-examined in the absence and presence of asciminib or dasatinib (see Supplementary Note 2 for additional details). In vehicle-treated cells from drug-treated mice, levels of pSTAT5 remained low indicating sustained inhibition of kinase signaling *in vivo* (MFI: 121±25, *p*=0.0125 asciminib-treated; MFI: 169±18, *p*=0.038 dasatinib-treated). Importantly, a further reduction in pSTAT5 was observed in cells from all cohorts upon asciminib or dasatinib treatment *ex vivo* indicating retained drug sensitivity (Figure 3B).

**Figure 3.**
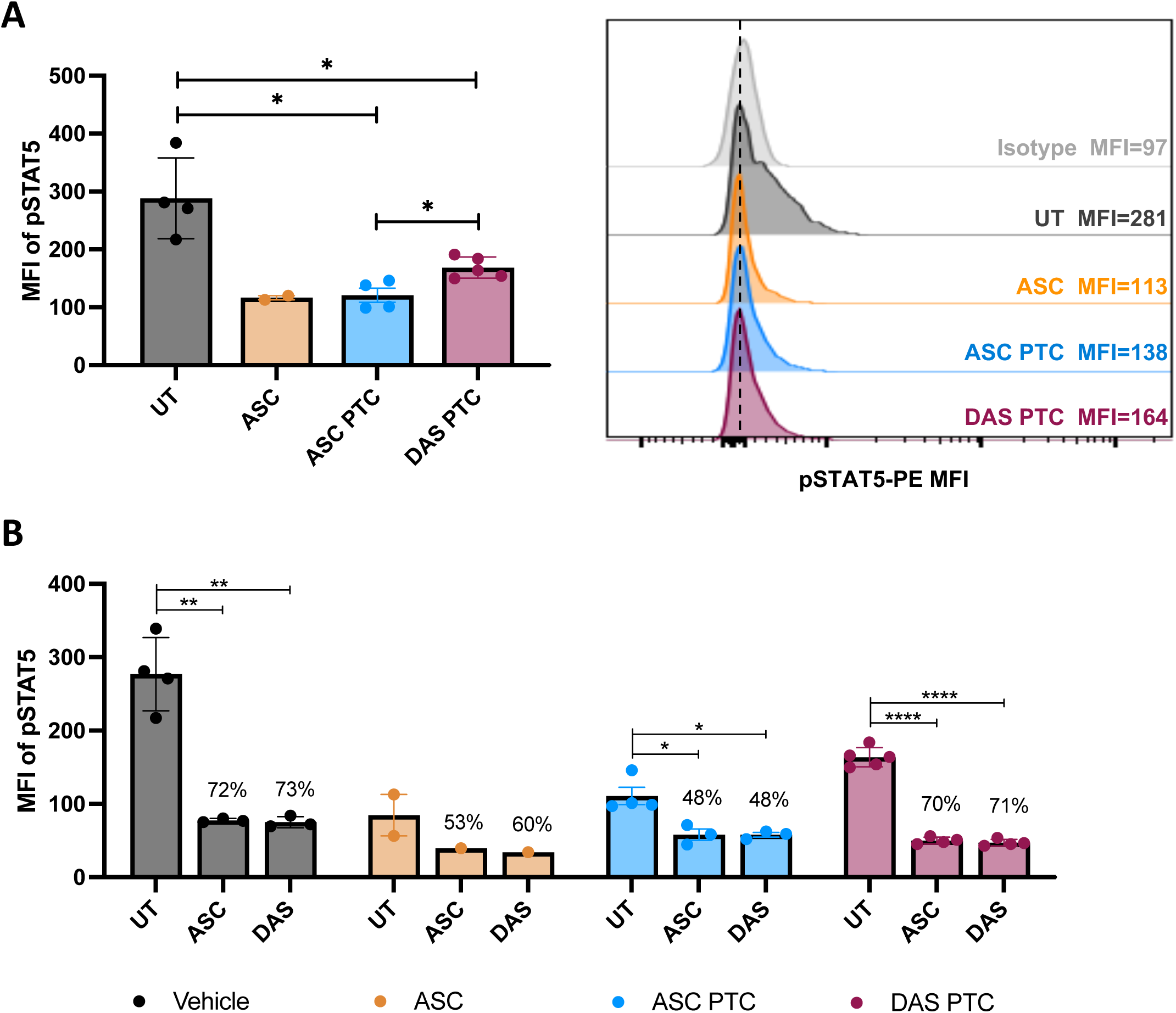
STAT5 activation is increased in vehicle-treated mice compared with asciminib-treated mice. Intracellular flow cytometry specific for pSTAT5 was performed *ex vivo* on cells harvested from *NUP214::ABL1* T-ALL PDX mice. A) Basal level of STAT5 activation in spleen cells harvested from vehicle-treated *NUP214::ABL1* mice compared with drug-treated mice. Representative flow cytometry histograms are shown on the right. B) For each mouse cohort, the basal level of STAT5 activation (UT, untreated) was investigated compared with cells treated with 5 µM asciminib or 100 nM dasatinib for >2 hours. pSTAT5 levels were normalized to isotype control. Percentage reductions in the presence of drug compared with untreated samples are specified. MFI=mean fluorescence intensity, UT=untreated, ASC=asciminib, DAS=dasatinib, PTC=post-treatment cessation. Individual data points are indicated, and Welch’s unpaired t-tests calculated significance where sample size allowed *p<0.05, ****p<0.0001.

### Deletion of specific regions of *ABL1* exon 3 confers varied response to asciminib in cells expressing *NUP214::ABL1*

Recent publications report the importance of the *ABL1* exon 2 encoded portion of the SH3 domain for protein folding and asciminib efficacy^48,49^. However, we now demonstrate *in vivo* efficacy of asciminib against *NUP214::ABL1* disease lacking *ABL1* exon 2. To investigate this further, Ba/F3 cells expressing full length *NUP214::ABL1* (e32a3 breakpoint) and *NUP214::ABL1* Δ1-4 isoforms (Figure S9; Supplementary Note 3) were assessed for ability to grow in the absence of IL-3. All cell lines were readily transformed to cytokine independence and retained kinase activity (Figure 4A). We then investigated kinase inhibition in the presence of asciminib versus dasatinib, and efficacy of both inhibitors in cell death assays. As expected, complete inhibition of STAT5 activity was observed in the presence of dasatinib (86-93% decrease depending on the Δisoform). Asciminib also inhibited kinase activity in cells expressing all four Δisoforms, but notably, the decrease in STAT5 activity for *NUP214::ABL1* Δ1 cells (49%) was significantly less than that observed for the other isoforms (Δ2=59%, *p*=0.0721; Δ3=66%, *p*=0.0342; Δ4=72%, *p*=0.0017; Figure 4B-F). Additionally, differing viabilities were observed upon treatment with asciminib. Ba/F3 cells expressing full-length *NUP214::ABL1* (LD_50_=158 nM) and *NUP214::ABL1* Δ2-4 (LD_50_=57, 43, 51 nM, respectively) demonstrated asciminib sensitivity within the clinically achievable range. Interestingly, *NUP214::ABL1* Δ1 cells were resistant to asciminib: LD_50_=18.3 μM, *p*<0.0001 compared with full-length *NUP214::ABL1* cells (Figure 5A). Dasatinib-mediated cell death was well within the clinically achievable range for all cell lines (LD_50_<5 nM) with no significant difference observed for any Δisoform compared with *NUP214::ABL1* full-length (*p*=0.4, Figure 5B). Importantly, Sanger sequencing of the entire *NUP214::ABL1* fusion gene confirmed the absence of any additional *ABL1* mutations in all cell lines (data not shown).

**Figure 4.**
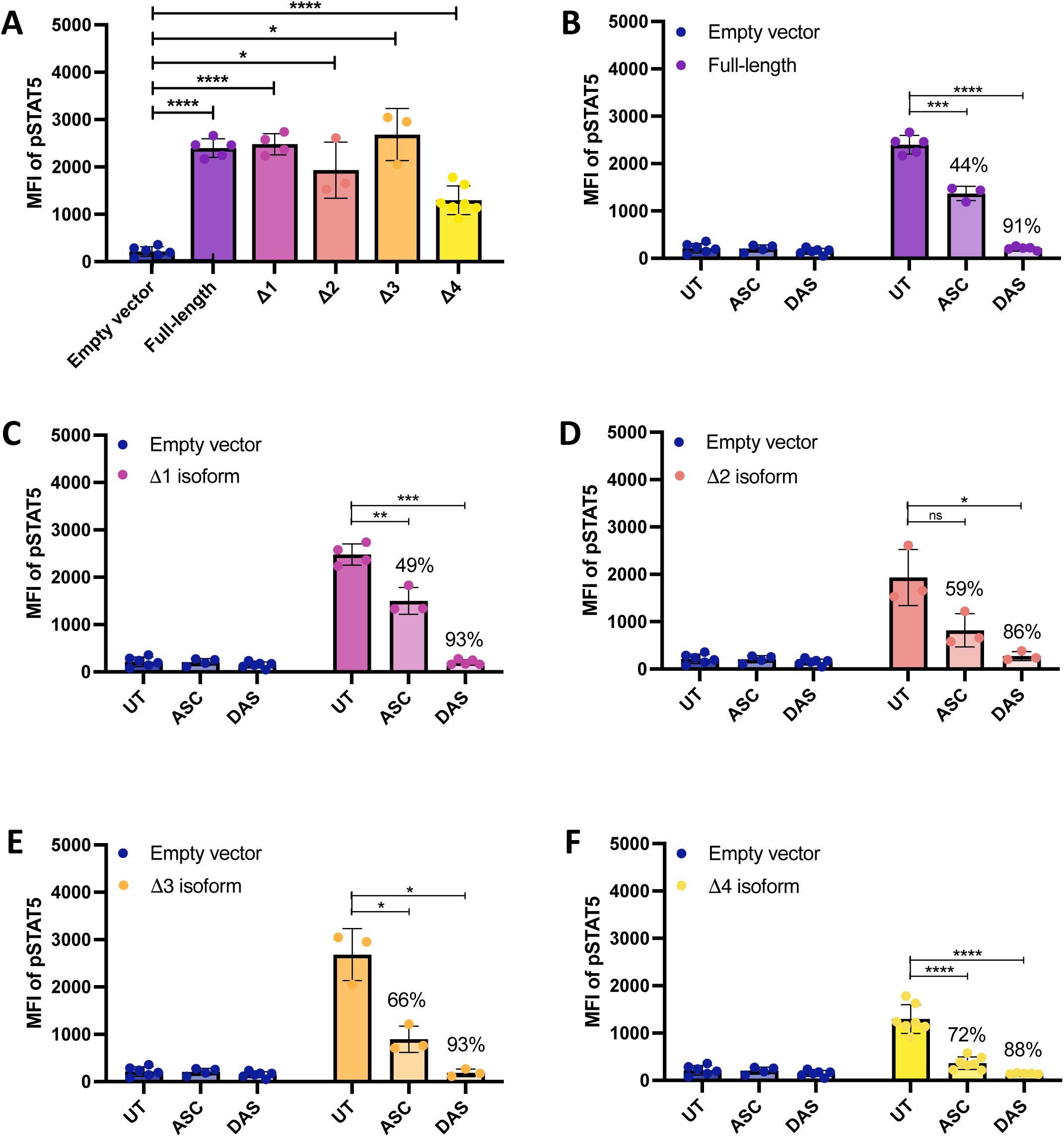
The constitutive activation of kinase signaling observed for all NUP214::ABL1 isoforms is reduced following treatment with asciminib. Signaling pathway activation of A) NUP214::ABL1 full-length and Δ1-4 isoforms was measured by intracellular flow cytometry specific for pSTAT5. B-F) inhibition of STAT5 activation in the presence of asciminib (5 µM) or dasatinib (100 nM) was also measured. Empty vector cells were washed in cytokine free media and rested for 4 hours before all cells underwent drug treatment for 2 hours. pSTAT5 levels were normalized to isotype control. A minimum of n=3 biological replicates were performed with individual data points indicated, and Welch’s unpaired t-tests calculated significance *p<0.05, **p<0.01, ***p<0.001, ****p<0.0001. Percentage reductions in the presence of drug compared with untreated samples are specified. MFI=mean fluorescence intensity, UT=untreated, ASC=asciminib, DAS=dasatinib.

**Figure 5.**
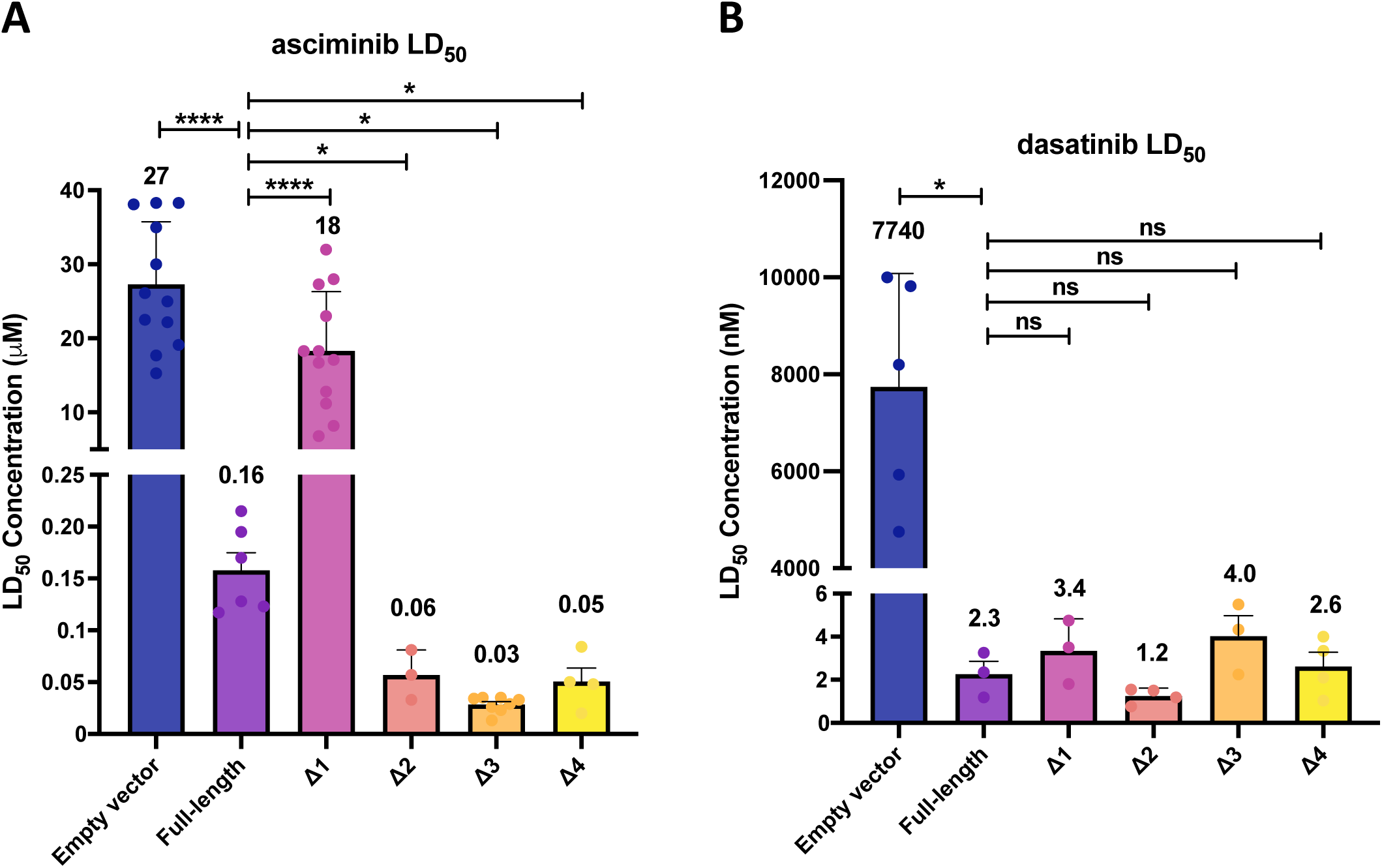
Asciminib demonstrates varying efficacy against NUP214::ABL1 isoforms *in vitro*. LD_50_ concentrations were calculated from cell death assays targeting different NUP214::ABL1 isoforms with A) asciminib B) dasatinib. The average LD_50_ concentrations written on the graphs are from a minimum of n=3 biological replicates with individual data points indicated, and Welch’s unpaired t-tests calculated significance *p<0.05, ****p<0.0001.

### Structural modelling of Δ1-4 isoform quaternary architecture

We next investigated the potential effect of the *NUP214* exon 32 fusion on ABL1 structure using modeling to derive a hypothesis for our *in vitro* findings. We generated structural models with AlphaFold3 for wildtype ABL1, NUP214::ABL1 (B-ALL Patient 2, e32a3 breakpoint), and NUP214::ABL1 Δ1-4 isoforms. The predicted structures showed conservation of the kinase domain fold, and both asciminib and nilotinib were successfully modeled into their respective binding sites (Figure 6A,C). Modeling supported binding of nilotinib and asciminib to NUP214::ABL1, as well as the Δ1-4 isoforms generated here, indicating alterations to the SH2 and SH3 domains are unlikely to significantly affect the structure of the inactive kinase domain.

**Figure 6.**
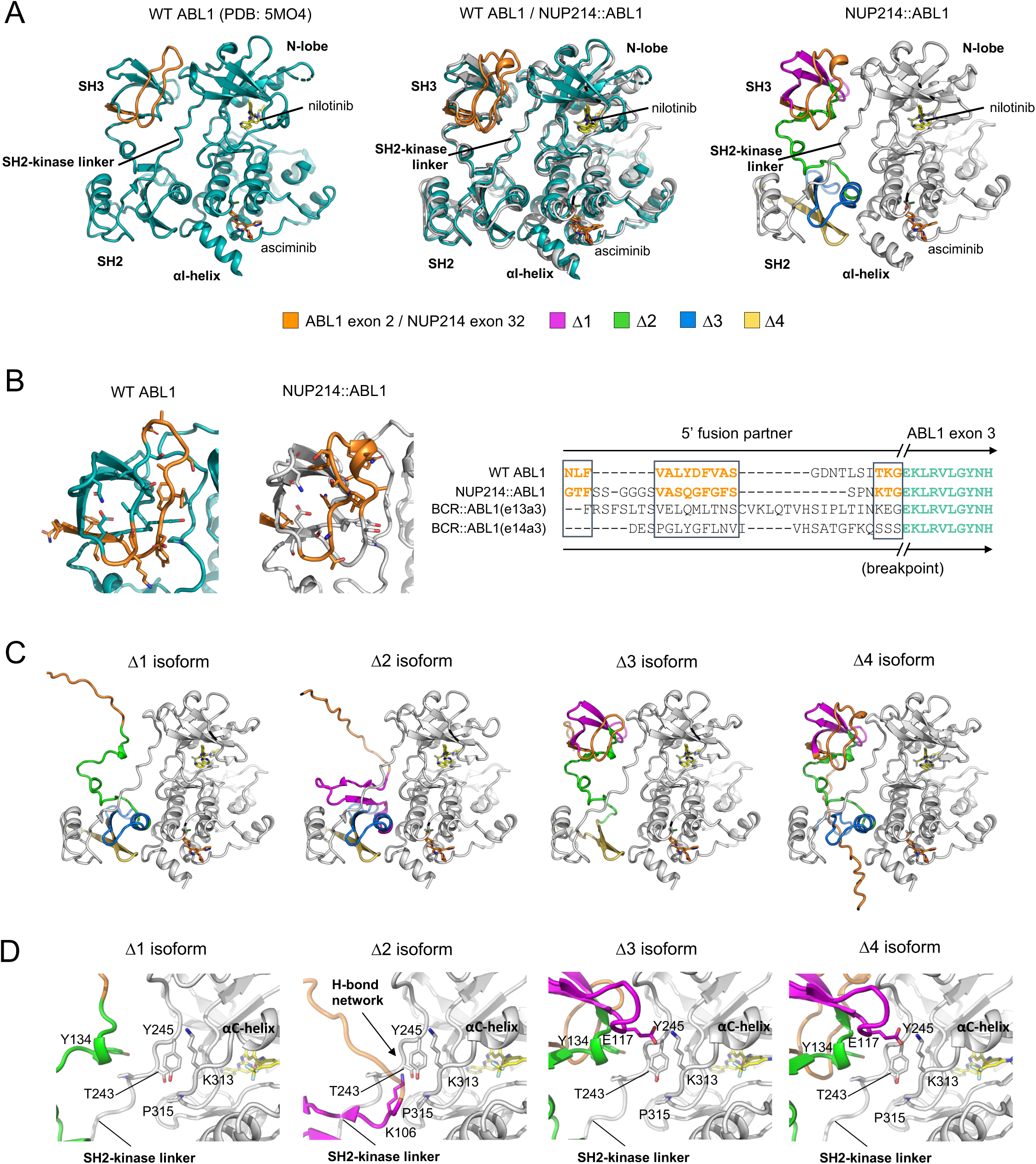
NUP214 exon 32 compensates for ABL1 exon 2 and is involved in formation of critical interactions between the ABL1 SH3 domain and N-lobe of the kinase domain. Representations of the crystal structures of A) ABL1 in its asciminib-bound autoinhibited conformation with exon 2 shown in orange (PDB: 5OM4); overlay of ABL1 (teal) and NUP214::ABL1 models (grey); and NUP214::ABL1 model with Δ1-4 regions depicted and NUP214 exon 32 shown in orange. B) left panel: a close-up view of the wildtype ABL1 SH3 domain compared with the same region of the NUP214::ABL1 fusion highlighting the functional compensation of NUP214 exon 32 maintaining protein folding and SH2-kinase linker and N-lobe interactions. Residue sidechains are shown as sticks. Right panel: sequence alignment comparing ABL1 exon 2, NUP214 exon 32, BCR exon 13 and BCR exon 14. The regions of sequence homology between ABL1 and NUP214 are delineated by boxes and coloured in orange in both panels. C) NUP214::ABL1 (e32a2) Δ1-4 isoforms highlighting the effects of each deletion region on the interactions between the SH3 domain, the SH2 domain, the kinase domain, and protein folding. D) A close-up view of each Δisoform highlighting formation of a salt bridge between p.E117 and p.K313 (Δ3 and Δ4), the absence of salt bridge formation (Δ1) and a new hydrogen bonding network between K106 (magenta) and the SH2-kinase linker in the Δ2 isoform (see arrow). In panels A, C and D, asciminib (orange) and nilotinib (yellow) are shown binding to the myristate and ATP-binding pockets, respectively.

Our *in vitro* data demonstrate that the NUP214::ABL1 cells (e32a3 breakpoint) remain sensitive to asciminib, which would require a functional SH3 domain to maintain allosteric inhibition^48,49,51^. Interestingly, the structural modeling indicates that a segment of NUP214 exon 32 may complete formation of the RT-loop, which could preserve protein function and interaction of the SH3 domain with the kinase domain (Figure 6B). Notably, exon 32 of NUP214 harbors sequence similarity with ABL1, which could rescue the loss of ABL1 exon 2 and maintain interactions with the SH2-kinase linker and N-lobe during protein folding. Indeed, equivalent sequences from *BCR::ABL1* e13a3 and e14a3 isoforms, which are asciminib resistant due to perturbation of the SH3 domain fold^48,49^, show a comparative absence of sequence homology (Figure 6B). Furthermore, when we compared the predicted folding of Δ1-4 isoforms in the setting of wildtype ABL1 versus the NUP214::ABL1 fusion, we found that Δ2-4 isoforms retained the SH3 fold when ABL1 exon 2 was present (as is the case in wildtype ABL1; Figure S10). However, in the absence of ABL1 exon 2 (as is the case in the NUP214::ABL1 fusion studied here), only Δ3 and Δ4 isoforms maintained SH3 folding (Figure 6C). This distinction highlights the importance of ABL1 exon 2 in maintaining protein folding and the compensatory role of the 5’ fusion partner (NUP214 exon 32).

### SH3 domain-mediated stabilization of the αC-helix is essential for asciminib efficacy

In wildtype ABL1, the SH3 domain interacts with the SH2-kinase linker and the kinase domain N-lobe, particularly the αC-helix, which are important for maintaining the auto-inhibited inactive conformation^41,42,53^. In the asciminib-resistant *NUP214::ABL1* Δ1 isoform, a section of the SH3 domain is deleted which abolishes a salt bridge between residues p.E117 (SH3 domain) and p.K313 (C-terminal end of the αC-helix; Figure 6D). SH3-mediated stabilization of the αC-helix is essential for auto-regulation and allosteric asciminib-dependent inhibition of kinase activity^54^. Mutation of p.K313 to glutamate confers asciminib resistance that is independent from asciminib effectively binding in the myristoyl pocket^43^. This salt bridge is predicted to be maintained in *NUP214::ABL1* Δ3-4 isoforms by proper folding of the SH3 domain, but not in the *NUP214::ABL1* Δ2 isoform. This should render *NUP214::ABL1* Δ2 cells resistant to asciminib, however, this is not the case (Figure 5A). Instead, in the Δ2 isoform, the SH3 β-sheet rests anti-parallel to the SH2 β-sheet and the SH2-kinase linker (Figure 6D). We hypothesize that this conformation would likely maintain stabilization of the αC-helix through stable docking against the SH2-kinase linker. Taken together, our findings show that complete deletion of the SH3 domain is not required for asciminib resistance, but rather only a small section that is essential for asciminib-mediated inhibition.

## Discussion

The STAMP compound asciminib, binds to a distinct and separate region of ABL from that where ATP-competitive TKIs bind, and can overcome resistance in patients previously treated with TKIs^39^ or can be used in combination with TKIs to restore treatment efficacy^43,55^. We have previously reported varying efficacy of asciminib against different *ABL1* and *ABL2* fusions identified in *ABL*r ALL^50^. In this study we explored potential reasons for these variations in efficacy using *in vitro* and *in silico* structural modeling. We also established three pre-clinical patient derived xenografts and confirmed *in vivo* asciminib efficacy against *NUP214::ABL1* disease in both T- and B-ALL settings.

Dasatinib is the current standard of care TKI used as an adjunct therapy for Ph+ ALL and *ABL*r ALL^29,32,56^. As dasatinib has demonstrated efficacy against *NUP214::ABL1* ALL *in vivo*^27,28^ it was used as comparator for successful treatment outcomes in our initial *NUP214::ABL1* T-ALL PDX model. Mice treated with asciminib as a single agent experienced similar survival to mice treated with dasatinib. In ALL three PDX models, asciminib treatment decreased splenomegaly, eradicated ALL blasts from the BM and liver, and significantly extended survival following treatment cessation. Our data provide a rationale for expansion of future asciminib clinical trials to include patients with Ph-like disease, particularly those with the *NUP214::ABL1* fusion.

There are 21 *ABL1* gene fusions reported in both B-ALL and T-ALL^16,57^, 19 of which have reported *ABL1* breakpoints in exon 2 (n=12), exon 3 (n=2) or exon 4 (n=8), plus 6 additional fusions where *ABL1* is the 5’ gene resulting in truncation of the kinase domain. Given that the majority of *ABL1* fusion genes harbor exon 2 or exon 3 breakpoints, and following our discovery that ABL1 fusions with exon 4 breakpoints are resistant to asciminib^50^, we used site-directed mutagenesis to interrogate the key region of ABL1 necessary for asciminib sensitivity. Four different *NUP214::ABL1* isoforms were created with large amino acid deletions within *ABL1* exon 3 partially encoding the SH3 domain; these isoforms also lacked *ABL1* exon 2 encoding the remainder of the SH3 domain (see Supplementary Note 3). Using cytokine-dependent Ba/F3 cells, we demonstrated each isoform retained the ability to induce cytokine-independent cell growth and constitutive activation of kinase signalling as measured by inhibition of STAT5 phosphorylation. All isoforms also retained varying levels of sensitivity to asciminib (reduction of pSTAT5) within the clinically achievable range. This was in contrast to two recent studies examining asciminib efficacy against different isoforms of BCR::ABL1 in the CML setting^48,49^. These studies reported cells harboring BCR::ABL1 isoforms lacking ABL1 exon 2 demonstrated constitutive STAT5 activation that was not reduced in the presence of asciminib. Authors postulated that *ABL1* exon 2 residues are essential for proper protein folding, including SH2 domain:myristoyl pocket interactions, with subsequent impacts on asciminib binding when BCR is the fusion partner. Conversely, our results demonstrate that asciminib is able to bind to NUP214::ABL1 fusions in the absence of the exon 2 residues and mediate kinase inhibition. However, asciminib is unable to induce apoptosis in cells lacking the initial part of ABL1 exon 3 (*NUP214::ABL1* Δ1 cells). The SH3-SH2-kinase domain interface is critical for auto-inhibition of ABL1^41,42,53,58^. Our results suggest that the Δ1 isoform negatively impacts binding of the SH3 domain to the SH2-kinase linker region (Figure 6D) which destabilizes the inactive conformation preventing asciminib-mediated cell death. Indeed, comparing all Δisoform cells, the smallest percentage decrease in pSTAT5 was observed in the *NUP214::ABL1* Δ1 cells (Figure 4) suggesting asciminib-mediated kinase inhibition is least effective in this cell line. Conversely, we hypothesize that the other Δisoforms, which delete regions of the SH2 domain, are allowing better binding of the SH3 domain to the SH2-kinase linker facilitating adoption of a more stable inactive conformation. This is supported by increased asciminib sensitivity in Δ2-4 cells compared with full length *NUP214::ABL1* cells and better inhibition of kinase signalling.

Structural comparison of the NUP214::ABL1 isoform model used in our *in vitro* experiments (e32a3 breakpoint that retains asciminib sensitivity) to wildtype ABL1 indicates predicted homology of the SH3 domain fold. This homology was not observed in asciminib-resistant BCR::ABL1 isoforms lacking ABL1 exon 2 and provides a putative basis for the discrepancy between our study and others investigating BCR::ABL1 fusions^48,49^. Interestingly, our modeling also showed that preservation of the SH3 domain did not occur for the Δ2 isoform, highlighting a region of ABL1 exon 3, that is likely important for mediating asciminib sensitivity (see Supplementary Note 4).

The structural models of asciminib-sensitive NUP214::ABL1 Δ3-4 isoforms predicted interaction between the partial SH3 domain (p.E117) and the C-terminus of the kinase domain αC-helix (p.K313) via a salt bridge, while Δ2 isoform allosterically anchors the αC-helix via the SH2-kinase linker. Allosteric modulation of the αC-helix has previously been demonstrated as critical for activation and regulation of kinases including ABL1^59,60^. Previous mutation of the residues identified here (thereby abolishing salt bridge formation) resulted in constitutive activation of the ABL1 kinase due to disruption of SH3-mediated stabilization of the αC-helix ^54^. Importantly, mutation of the αC-helix lysine to glutamate directly confers asciminib resistance^43^. Deletion of the Δ1 region incorporating the p.E117 residue abolishes salt bridge formation and would prevent SH3-mediated stabilization of the kinase domain.

We report that asciminib sensitivity is retained in Δ2-4 cells harboring deletions within the SH2 domain. This contrasts with observations in the context of BCR::ABL1 where a degree of resistance occurs upon total deletion of the SH2 domain^49^. Our structural models show that the modified SH2 domain retains proximity to the myristoyl binding pocket and maintains stabilizing interactions with the SH2-kinase linker, potentially through salt bridge formation (observed for Δ3-4 isoforms), *de novo* interactions with NUP214 exon 32 (observed for Δ2 isoform) or other molecular interactions occurring between the SH3 domain, SH2-kinase linker and the N-lobe. Our findings reveal a previously undiscovered adaptability in allosteric regulation of ABL1 kinase activity by asciminib which may have important considerations for potential allosteric resistance mechanisms as well as future therapeutic targets and drug design.

The recent approval of asciminib as a first-line treatment for CML, combined with its excellent safety and tolerability profile, make it an attractive addition to the treatment repertoire for Ph+ ALL. We now report for the first time, pre-clinical drug studies that establish asciminib as a potential treatment for ALL driven by the *NUP214::ABL1* gene fusion. Our findings also identify a key region of the ABL1 SH3 domain critical for asciminib activity against NUP214::ABL1. These data significantly contribute to our understanding of asciminib’s mode of action, further characterize which ABLr ALL patients would benefit asciminib therapy and provide an additional therapeutic option for patients who are refractory to ATP-competitive TKIs or who develop resistance. The results presented here strongly support clinical trials evaluating asciminib efficacy in all *NUP214::ABL1* patients regardless of ABL1 breakpoint location.

## Methods

### ALL patient samples

We studied leukemia samples from biobanked material from three cases with *NUP214::ABL1* disease identified by mRNA-Seq, one T-ALL relapse sample and two B-ALL diagnosis samples. Details for patient samples are provided in Table S1 and Figure S1. Patients gave written informed consent for sample collection and storage for future ethically approved research including the studies presented here. All research was carried out in accordance with the Declaration of Helsinki.

### Inhibitor preparation

For *in vitro* experiments dasatinib and asciminib were obtained from Selleckchem (Houston, TX) and dissolved in DMSO (Sigma-Aldrich, St. Louis, MO) to a concentration of 10 mM and stored at 4°C. For *in vivo* experiments, dasatinib (Selleckchem) was dissolved in 80 mM citric acid (pH 3.1) and asciminib (free base solution form, a kind gift from Novartis Pharma AG; Basel, Switzerland) was suspended in 30% PEG300, 6% Solutol HS15, 1 M NaOH, 0.1 M HCl, 10 mM sodium acetate buffer (pH 4.7) strictly adhering to Novartis instructions. Dosing solutions were freshly prepared weekly.

### Development of ALL patient derived xenografts (PDX) and *in vivo* drug treatments

Six to 12 week old NOD.Cg-Prkdc^scid^Il2rg^tm1Wjl^/SzJ (NSG) mice (The Jackson Laboratory, Bar Harbor, ME) were treated subcutaneously with 0.1 mg Baytril in 0.9% sodium chloride per 10 g bodyweight prior to sub-lethal gamma-irradiation at 200 cGy. Spleen- or bone marrow (BM)-derived ALL patient blasts isolated from primary xenografts (1.5-1.8⊆10^6^ cells) were injected into the tail vein of secondary recipients. Engraftment and disease progression was monitored by weekly blood sampling from the tail vein and flow cytometric analysis of hCD45+ cells and hCD7+ (T-ALL PDX) or hCD19+ (B-ALL PDXs). Once hCD45+ levels reached a median of 5%, mice were randomized into treatment groups (n=5) as indicated A) vehicle control via oral gavage (For the T-ALL PDX: citric acid; for the B-ALL PDXs: HCl, PEG300, Solutol HS15, NaOH, sodium acetate solution) B) 20 mg/kg/day once daily dasatinib via oral gavage (T-ALL PDX only) C) 30 mg/kg once daily asciminib via oral gavage. Treatment was for 28 days and mice were maintained on 0.3 mg/mL enrofloxacin supplemented water for the duration of the experiment. Animals were monitored daily and were humanely killed when displaying either clinical signs of leukemia (eg: weight loss, reduced movement, ruffled fur) or when blood levels of hCD45+ cells were >50%. At experimental end, cardiac bleed and complete blood count (CBC) where possible were performed, and spleen, liver, thymus (where evident) and BM were harvested. Flow cytometric immunophenotyping was performed on peripheral blood and/or BM and peripheral organs, and formalin fixed organ sections were stained with hematoxylin and eosin (H&E) by the University of Adelaide Medical School Histology Service and scanned by the SAHMRI Light Microscopy and Laser Capture Core Facility. Slides were visualized with CaseViewer software (v2.2, 3DHISTECH^TM^ Ltd.). All experiments were performed on protocols approved by the SAHMRI Animal Ethics Committee.

### PCR validation of *NUP214::ABL1* breakpoint in PDX samples

Following extraction of RNA from PDX samples, cDNA was synthesized and PCR to verify fusion expression was performed with Platinum™ SuperFi II DNA Polymerase (ThermoFisher Scientific, Waltham, MA) according to the manufacturer’s instructions. Primer sequences for RT-PCR of fusion breakpoints and PCR conditions are detailed in the tables below.

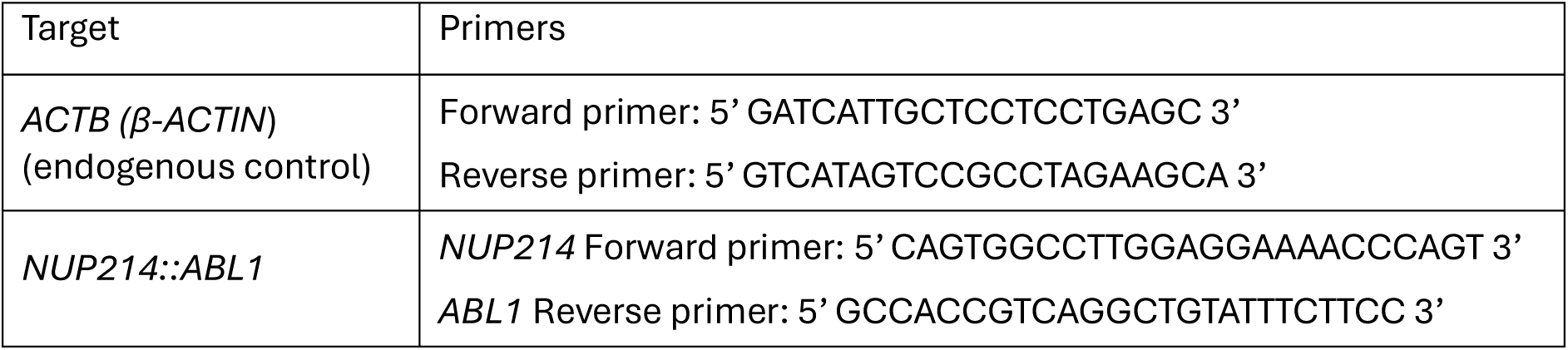

PCR cycling conditions for amplification of fusion breakpoints:

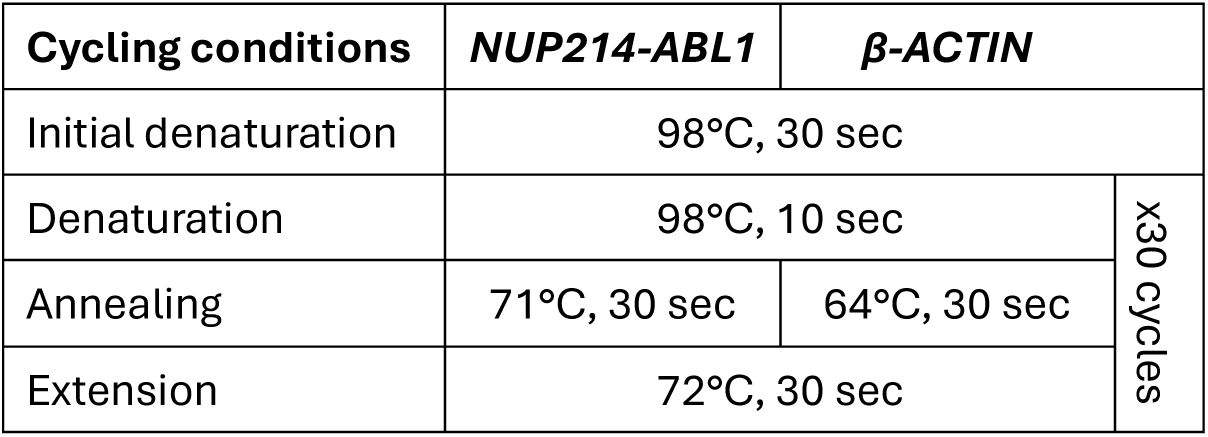
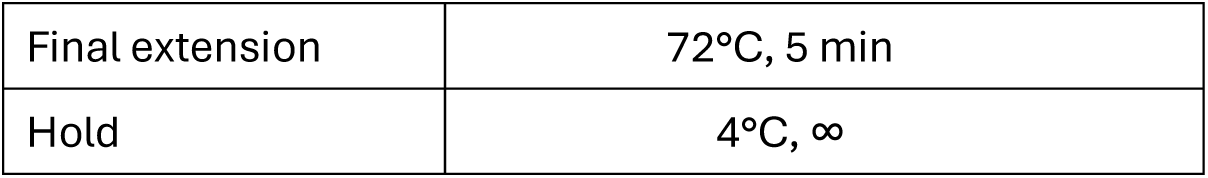

| Cycling conditions | NUP214-ABL1 | β-ACTIN | x30 cycles |
| --- | --- | --- | --- |
| Initial denaturation | 98°C, 30 sec |  |  |
| Denaturation | 98°C, 10 sec |  |  |
| Annealing | 71°C, 30 sec | 64°C, 30 sec |  |
| Extension | 72°C, 30 sec |  |  |

|  |  |
| --- | --- |
| Final extension | 72°C, 5 min |
| Hold | 4°C, ∞ |

### Site directed mutagenesis of *NUP214*::*ABL1*

The *NUP214::ABL1* fusion gene was amplified from the cDNA of a B-ALL patient with a *NUP214*::*ABL1* (e32a3) fusion (B-ALL Patient 2) and subcloned into a Gateway pDONR-221 vector (Invitrogen, Carlsbad, CA) using the Gateway BP Clonase II enzyme mix (Invitrogen) as described previously^50^. We designed four separate retroviral constructs that deleted regions of ∼25 amino acids each from *ABL1* exon 3 (Figure S9, Table S2) by site directed mutagenesis (SDM) and designated these constructs Δ1-4. The rationale for deleting this region is provided in Supplementary Note 3. *NUP214::ABL1* full length served as control. SDM primers were designed using NEBaseChanger software (New England Biolabs [NEB], Ipswich, MA) and PCR conditions were specific for each primer pair (detailed in the table below). Following Sanger sequencing confirmation of deletion of the correct sequence, *NUP214::ABL1* Δ1-4 fusion genes were then transferred into a Gateway-compatible pMSCV-IRES-GFP (pMIG) vector using the Gateway LR Clonase II enzyme mix as per the manufacturer’s instructions (Invitrogen) before generation of viral supernatants and transformation of Ba/F3 cells.

Optimized cycling conditions for site-directed mutagenesis:

| Cycling conditions | Δ1 | Δ2 | Δ3 | Δ4 |
| --- | --- | --- | --- | --- |
| Initial denaturation | 98°C, 30 sec |  |  |  |
| Denaturation | 98°C, 10 sec |  |  |  |
| Annealing | 63°C, 30 sec | 69°C, 30 sec | 72°C, 30 sec | 68°C, 30 sec |
| Extension | 72°C, 8 min |  |  |  |
| Final extension | 72°C, 2 min |  |  |  |
| Hold | 4°C, ∞ |  |  |  |

### Intracellular flow cytometry

Following a 4 h cytokine starvation (Ba/F3 empty vector cells only) and 2 h drug treatment, Ba/F3 cells and thawed leukemic blasts from patient derived xenografts (PDX) were fixed with a final concentration of 1.6% paraformaldehyde (ThermoFisher Scientific) for 10 min, washed in 1⊆PBS and permeabilized with 80% methanol overnight at -80°C. Cells were then washed in 1⊆PBS, and subsequently in 1⊆PBS/1% bovine serum albumin (BSA, Sigma Aldrich). Cells were stained with antibodies targeting pSTAT5 (BD Biosciences) or IgG1 control (BD Biosciences and DakoCytomation) and all intracellular staining was carried out in the dark for 60 min at room temperature in 1⊆PBS/1% BSA. Cells were washed in 1⊆PBS before data collection on a BD FACSCanto™ analyzer.

### *Escherichia coli (E.coli)* transformation and plasmid purification

SDM products were combined with KLD reaction buffer and KLD enzyme mix (NEB) according to the manufacturer’s instructions. Following incubation at room temperature for 60 min, 5 μL KLD reaction was combined with DH5α *E. coli* competent cells before further incubation for 30 min on ice. Samples were then heat shocked at 42°C for 30 sec, incubated on ice for 5 min, then combined with S.O.C media (Invitrogen) and incubated at 37°C for 90 min with agitation. Plasmid-containing *E.coli* cells underwent antibiotic selection (kanamycin for pDONR, ampicillin for p-MIG) before expansion in antibiotic-containing Luria Broth. Plasmid DNA was purified using QIAprep Spin Miniprep Kit (Qiagen, Hilden, Germany] as per manufacturer’s instructions before quantitation using a Qubit^®^ 2.0 fluorometer (Invitrogen).

### Viral Transduction

Retrovirus was produced by transfecting 1⊆10^6^ HEK293T cells in 5 mL recipient cell media in a T25 culture flask with 4 µg of each *NUP214::ABL1* expression vector, 4 µg of the pEQ-Eco packaging vector and 20 µL lipofectamine (Invitrogen). Viral supernatant was harvested 48 h post transfection, centrifuged and passed through a 0.45 µm filter. Ba/F3 cells at a concentration of 3⊆10^5^ cells/mL were centrifuged at 1800 rpm for 1 h with 30 µg/mL polybrene (Santa Cruz Biotechnology, Dallas, TX) in 4 mL of viral supernatant in a 6-well plate at room temperature. Cells were washed 24 h later and sub-cultured in original media. Ba/F3 cells were sorted at a concentration of 1⊆10^7^ cells/mL in RPMI and 2% FCS on a BD FACSAria™ (BD Biosciences, Franklin Lakes, NJ) for GFP at >95% purity.

### Cell lines and maintenance

HEK293T (ATCC, Manassas, VA) cells were maintained in DMEM (Sigma-Aldrich, St. Louis, MO) supplemented with 10% Fetal Calf Serum (FCS; CellSera Australia, Rutherford, NSW). Ba/F3 cells (ATCC) were maintained in RPMI (Sigma-Aldrich) supplemented with 5% WEHI-3B conditioned media as a source of murine IL-3 for Empty vector control cells^61^. Cells expressing full length and *NUP214::ABL1* Δ1-4 constructs were cultured in cytokine free media. All cell line media contained 10% FCS, 200 mM L-Glutamine (SAFC Biosciences, Lenexa, KS), 5000 U/mL penicillin (Sigma-Aldrich) and 5000 µg/mL streptomycin sulphate (Sigma-Aldrich). All cell lines were mycoplasma negative according to MycoAlert™ Mycoplasma Detection Kit (Lonza, Basel, Switzerland).

### Annexin V/7-AAD cell death assay

Ba/F3 cell death was assessed by seeding at 3.5⊆10^4^ cells/mL in a 24-well plate in the presence of a drug concentration gradient. Empty vector control cells were plated in the presence of 0.5% IL-3 conditioned supernatant, while *NUP214::ABL1* expressing cells were plated in cytokine free media for a 3-day cell death assay. At 72 h, cells were resuspended, plated in a 96-well plate and stained with 0.4 µL AnnexinV-PE (BD Biosciences) and 0.04 µL 7-AAD (ThermoFisher Scientific) in 20 µL Hank’s Balanced Salt Solution (HBSS, Sigma Aldrich) with 1% HEPES solution (Sigma Aldrich) and 5% CaCl_2_ for 20 min on ice in the dark. Cells were washed in 1⊆PBS before data collection on a BD LSFortessa™ analyzer (BD Biosciences).

### Structural modeling and molecular docking

Structural models were predicted for full-length *NUP214::ABL1* and each Δ1-4 isoform, as well as wildtype ABL1 using AlphaFold3^52^. To model the binding modes of asciminib and nilotinib, each compound was docked to each structure in ICM-Pro (Molsoft L.L.C, San Diego, CA) and rigorously energy minimized; each complex was then refined. The models shown in the main text were selected based on similarity of each domain to the crystal structure of ABL1 (5MO4) and clear residue interactions. An alignment of each of the five models output from AlphaFold3 (coloured by pLDDT score) are provided in Figures S12-13. Structural analysis and visualization was performed in PyMOL (Schrodinger, INC) and a detailed interpretation of the results is provided in Supplementary Note 4. Primary amino acid sequences of ABL1 and NUP214 were obtained from UniProt under the accession numbers P00519 (ABL1) and P35658 (NUP214).

### Quantification and statistical analyses

GraphPad Prism software version 10.3.0 (GraphPad Software Inc.), FlowJo software version 10.6.1 (FlowJo LLC) were used for analyses. All assays were carried out in triplicate and graphs represent the median value or mean with standard deviation (SD) error bars as indicated in the figure legends. Unpaired t-test or one way ANOVA was used to determine the difference between experimental groups. Kaplan-Meier survival curves were analyzed using a log-rank test. Differences were considered statistically significant when the *p*-value was <0.05. *p<0.05, **p<0.01, ***p<0.001.

### Ethics Statement

Cryopreserved samples from patients diagnosed with Ph-like ALL within established biobanks were used in this study. The samples were obtained with informed consent for future unspecified research and to perform laboratory assays on existing samples. All patients were required to provide informed consent for the use of their samples in accordance with the Declaration of Helsinki. The nature of the study was described, and the required patient information signed consent (PISC) forms were issued to all patients. The forms described the questions that would be relevant to their involvement and when this was signed, gave consent for inclusion of the patient. The use of the samples was approved by the Women’s and Children’s Health Network Human Research Ethics Committee.

## Acknowledgements

Authors would like to thank Dr Andreas Bruederle (Global Biomarker Diagnostic Director, Novartis) for insightful and valuable comments. This research was undertaken with the following financial and other support to L.N.E.: the Cancer Council SA’s Peter Nelson Leukaemia Research Fellowship on behalf of its donors (ID# 022212); and the Cancer Council SA’s Beat Cancer Project on behalf of its donors and the State Government through the Department of Health (ID# MCR2257). The project was also undertaken with the financial support of the SAHMRI Early and Mid-Career Researcher Seed Funding Grant. Flow cytometry analyses and FACS were performed at the SAHMRI ACRF Cellular Imaging and Cytometry Core Facility. The Facility is generously supported by the Detmold Hoopman Group, the Australian Cancer Research Foundation, and the Australian Government through the Zero Childhood Cancer Program. Animal models were performed in the SAHMRI Bioresources Core Facility and we would like to acknowledge the technical support provided. Pediatric patient samples were provided by the Queensland Children’s Tumour Bank, which is generously supported by the Children’s Hospital Foundation (Queensland).

## Author Contributions

L.N.E. designed the study, performed *in vitro* and *in vivo* experiments, analyzed the data, wrote the manuscript, and created the figures; D.P.M. performed *in silico* modelling, provided scientific insight, wrote the manuscript, and created the figures; E.L. performed *in vitro* experiments, analyzed the data, provided scientific insight and drafted the manuscript; C.E.S. performed *in vitro* and *in vivo* experiments, and analyzed the data; T.S.C. performed *in vitro* experiments, and analyzed the data; E.C.P. performed *in vitro* experiments, analyzed the data, and drafted the manuscript; A.S.M. provided patient material for the PDX models; D.T.Y. provided patient material for the PDX models and scientific insight; T.P.H. designed the study, and provided scientific insight; D.L.W. designed the study, and provided scientific insight; and all authors critically reviewed the manuscript.

