## Supplementary Figures for "The 5’ fusion partner modulates response to the STAMP inhibitor asciminib in *ABL*-rearranged ALL"

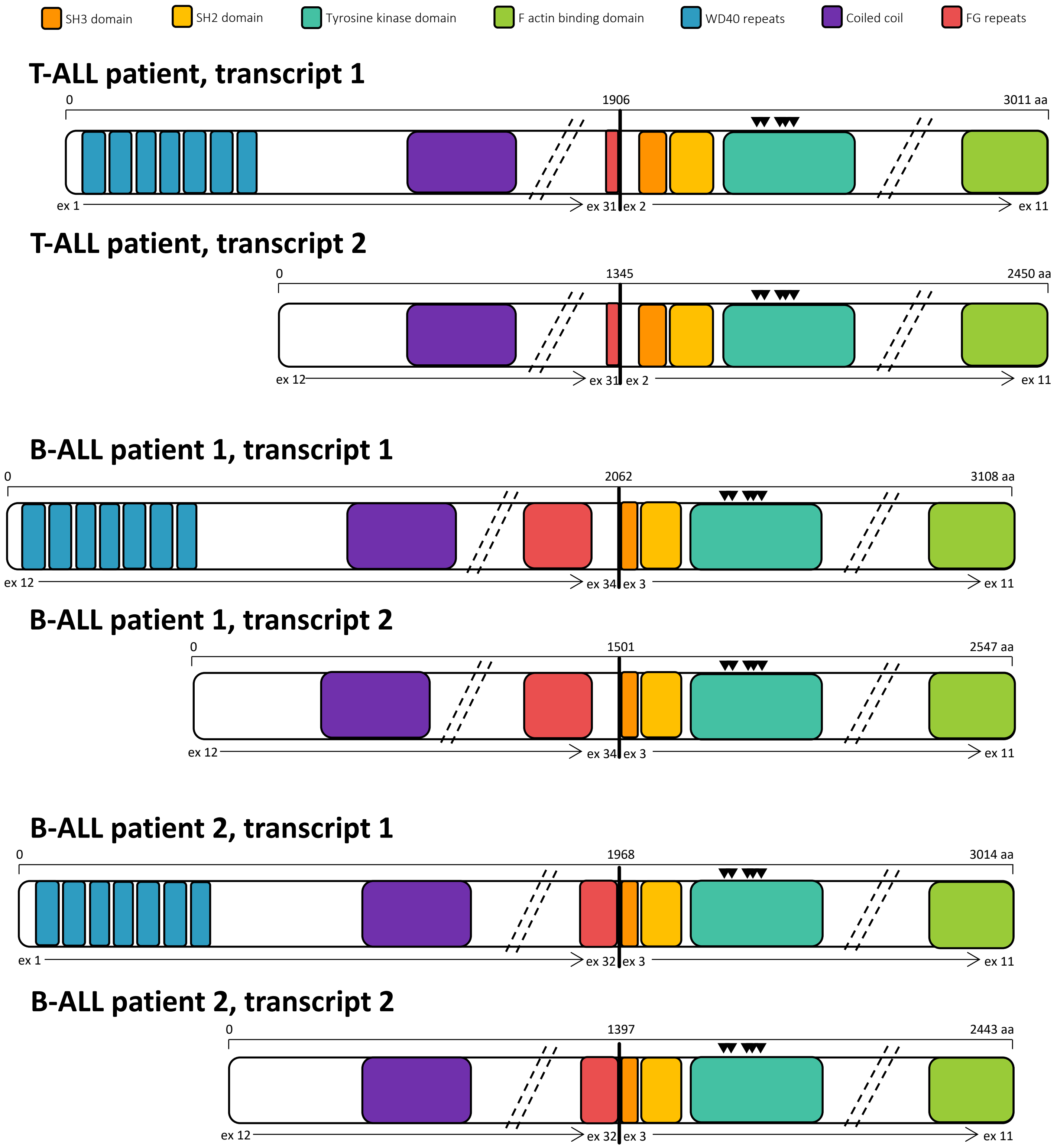


**Supplementary Figure 1. Schematic representation of *NUP214::ABL1* transcripts identified in one T-ALL patient and two B-ALL patients.** mRNA-sequencing identified two *NUP214::ABL1* transcripts In each of the patients due to two ATG codons. For the T-ALL patient, the 5' *NUP214* partner gene is fused to *ABL1* exons 2-11 retaining the complete ABL1 SH3 domain while for both B-ALL patients *NUP214* is fused to *ABL1* exons 3-11 truncating part of the ABL1 SH3 domain. Transcript 2 from B-ALL patient 2 was amplified for *in vitro* experiments. In all isoforms, the residues required for formation of the myristate binding pocket are retained indicated by black triangles above the kinase domain (▼). Breakpoints are denoted by vertical bold black lines and exons by vertical dashed lines. Diagonal dashed lines indicate unstructured regions of genes that have been omitted for simplicity. The *NUP214* NM_005085 and *ABL1* NM_007313 isoforms were used for schematic construction.

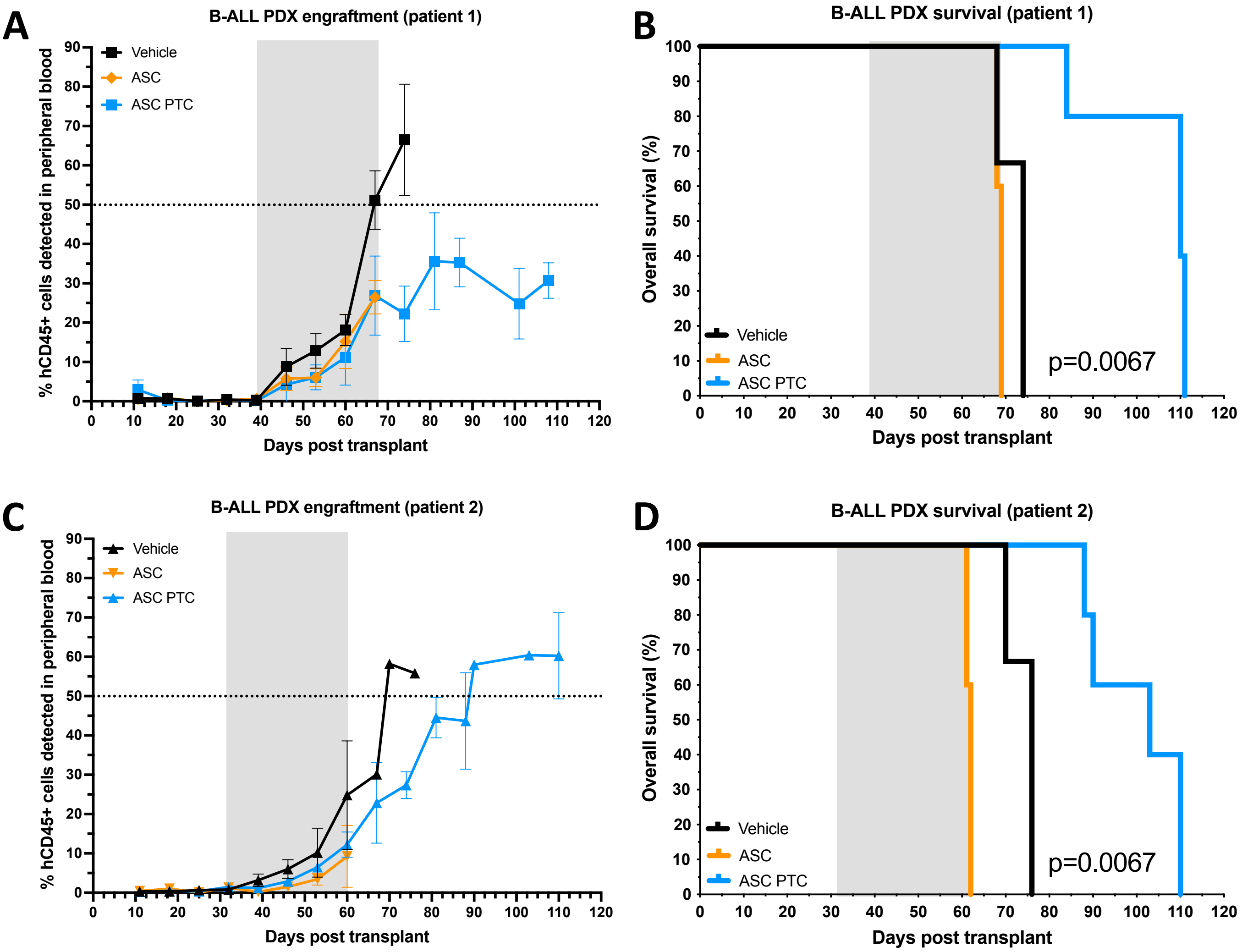


**Supplementary Figure 2. Treatment with asciminib significantly increases survival in two B-ALL *NUP214::ABL1* mouse models.** Patient derived xenografts (PDX) were established from two B-ALL patients. A,C) Engraftment of leukemic cells was tracked via measurement of hCD45+ cells in the peripheral blood. Points on the graphs represent mean ±SD. Treatment windows are highlighted in grey. B,D) Kaplan-Meier curves determined survival. Significance was evaluated between all curves and determined by Log-rank test. ASC=asciminib, PTC=post-treatment cessation. **NOTE**: B-ALL Patient 1 mice from the ASC PTC cohort were humanely killed prior to detection of 50% hCD45+ due to clinical signs of disease.

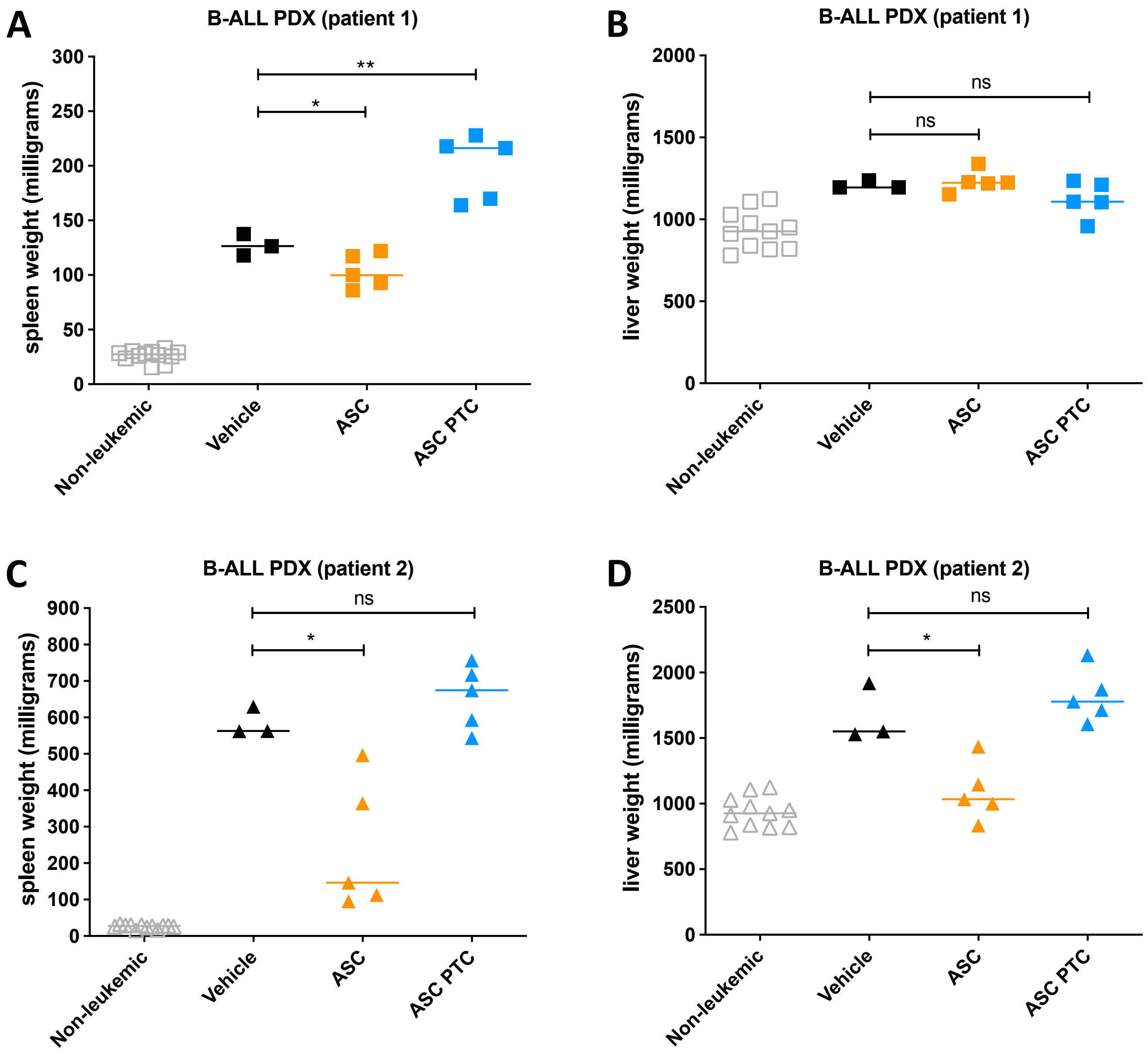


**Supplementary Figure 3. Treatment with asciminib reduces spleen and liver weights in two B-ALL *NUP214::ABL1* mouse models.** Patient derived xenografts (PDX) were established from two B-ALL patients. A,C) Spleen or B,D) liver weights of mice at experimental endpoint, either on treatment or post-treatment cessation. Values represent mean ± SD, horizontal lines represent the median. Welch's unpaired t-test was used to determine significance *p<0.05, **p<0.01. DAS=dasatinib, ASC=asciminib, PTC=post-treatment cessation.

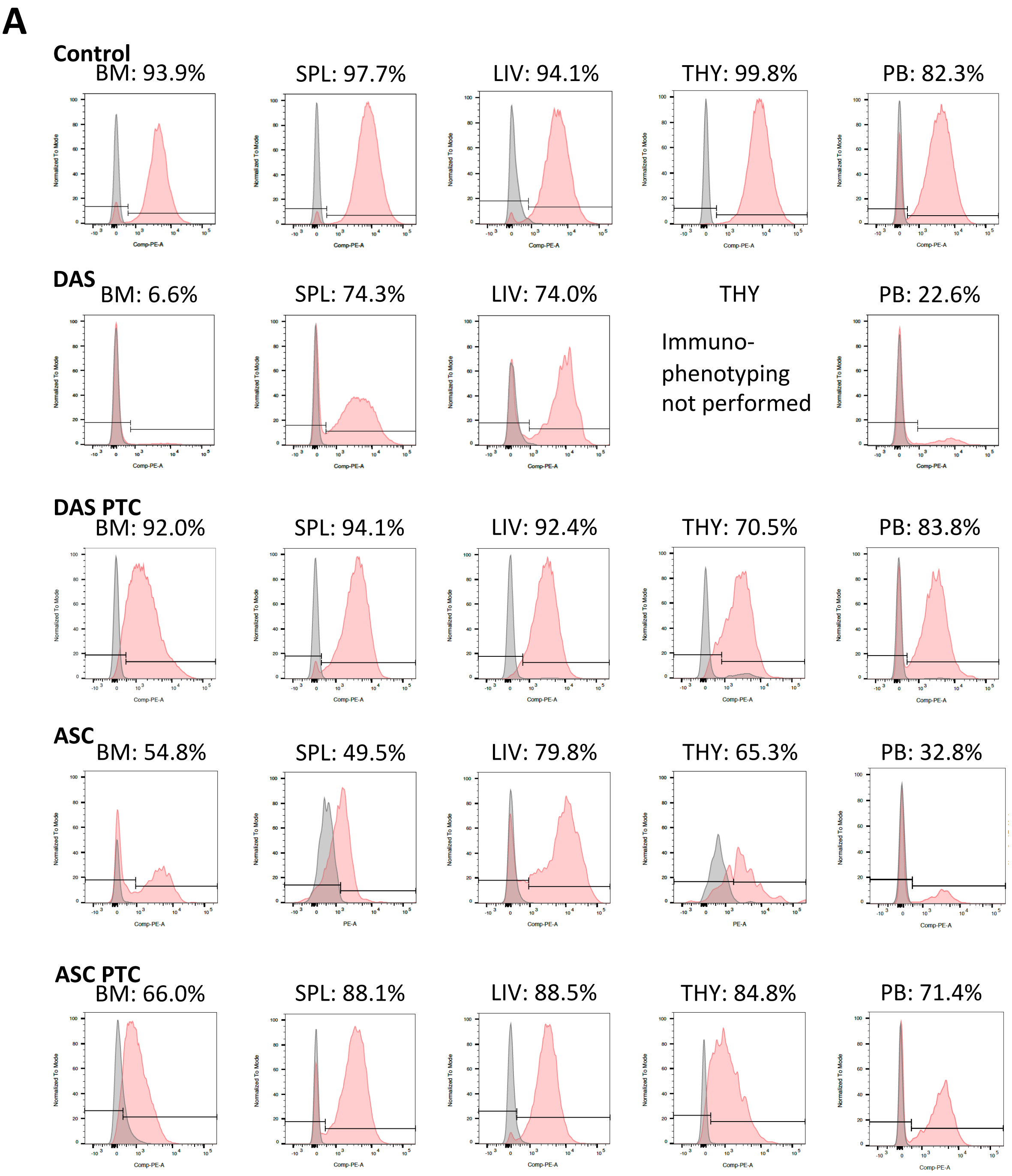


**Supplementary Figure 4. Characterization of leukemic infiltration in *NUP214::ABL1* T-and B-ALL PDX mice at experimental end point, either on treatment or post-treatment cessation. A)** Representative flow cytometric analyses for T-ALL PDX leukemic blasts stained with hCD7-PE antibody **B)** B-ALL Patient 1 and **C)** B-ALL Patient 2 PDX leukemic blasts stained with hCD19-PECy7 antibody at experimental end point. Cells were compared with isotype controls and percentage positivity for human cells calculated. BM=bone marrow, SPL=spleen, LIV=liver, THY=thymus, PB=peripheral blood.

**B**

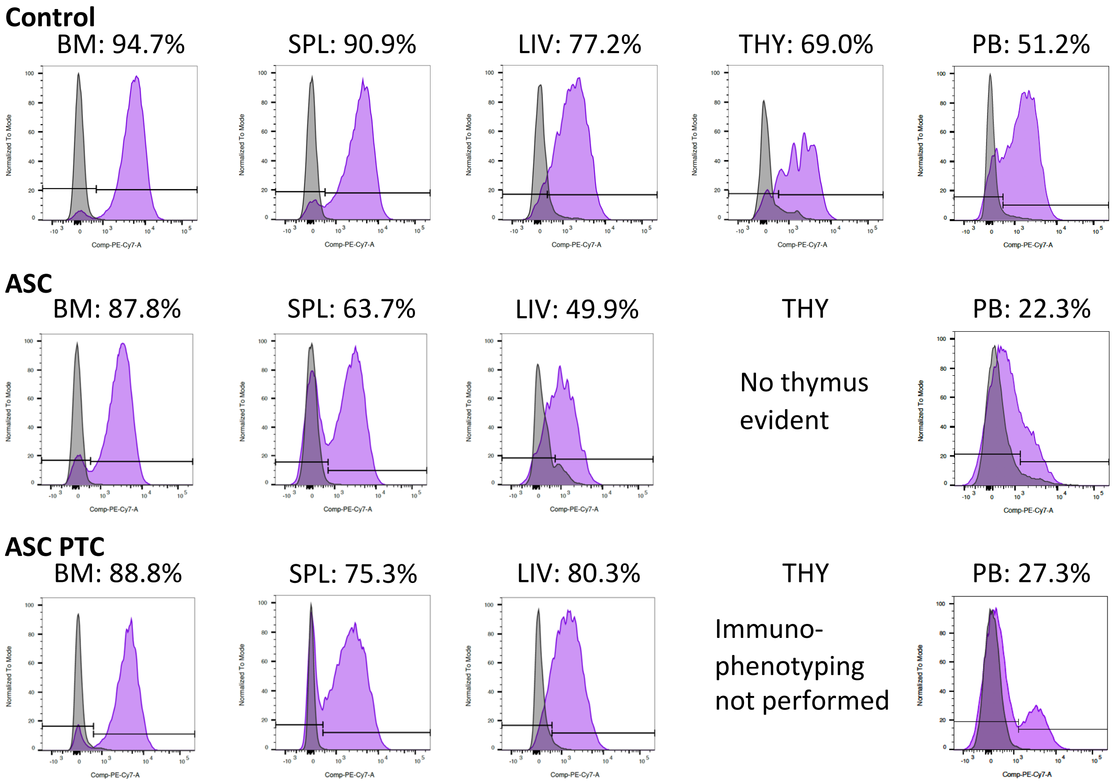
Control
BM: 94.7%
SPL: 90.9%
LIV: 77.2%
THY: 69.0%
PB: 51.2%
ASC
BM: 87.8%
SPL: 63.7%
LIV: 49.9%
THY
No thymus evident
PB: 22.3%
ASC PTC
BM: 88.8%
SPL: 75.3%
LIV: 80.3%
THY
Immuno-phenotyping not performed
PB: 27.3%
Normalized To Mode
Comp-PE-Cy7-A

**C**

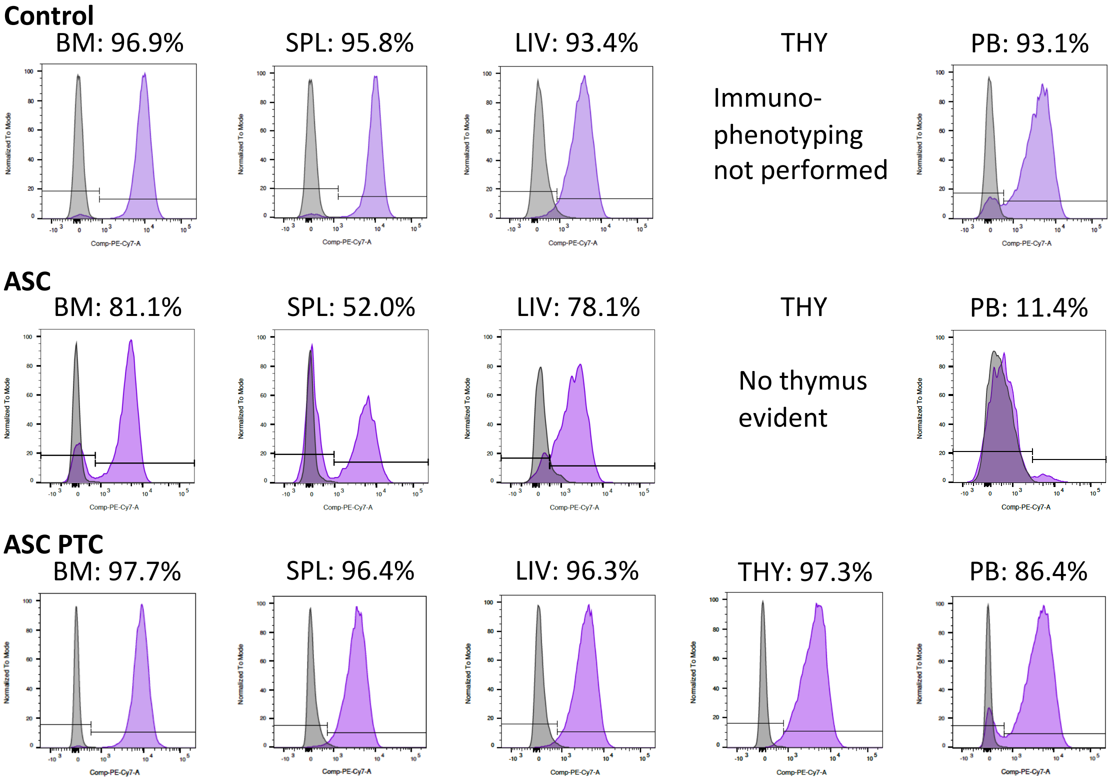
Control
BM: 96.9%
SPL: 95.8%
LIV: 93.4%
THY
Immuno-phenotyping not performed
PB: 93.1%
ASC
BM: 81.1%
SPL: 52.0%
LIV: 78.1%
THY
No thymus evident
PB: 11.4%
ASC PTC
BM: 97.7%
SPL: 96.4%
LIV: 96.3%
THY: 97.3%
PB: 86.4%
Normalized To Mode
Comp-PE-Cy7-A

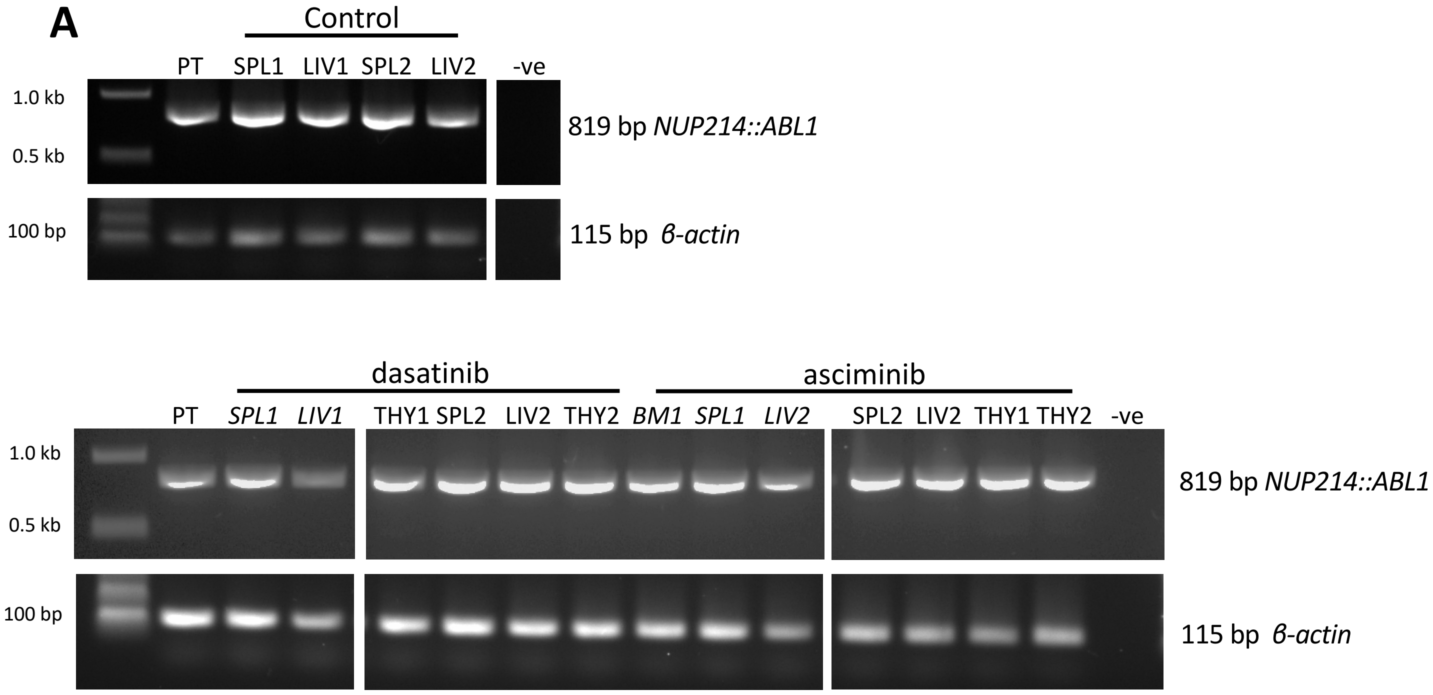


**Supplementary Figure 5. Characterization of leukemic infiltration in *NUP214::ABL1* T-ALL PDX mice at experimental end point, either on treatment or post-treatment cessation.** *NUP214::ABL1* breakpoint RT-PCR to confirm expression of *NUP214::ABL1* in **A)** T-ALL **B)** B-ALL Patient 1 **C)** B-ALL Patient 2 PDX samples. PCR products were separated by agarose gel electrophoresis and products detected at the expected sizes as indicated. On-treatment samples are delineated by italics. The NEB 1kb ladder was used to determine product size. PT=patient positive control, BM=bone marrow, SPL=spleen, LIV=liver, THY=thymus, -ve=water negative control.

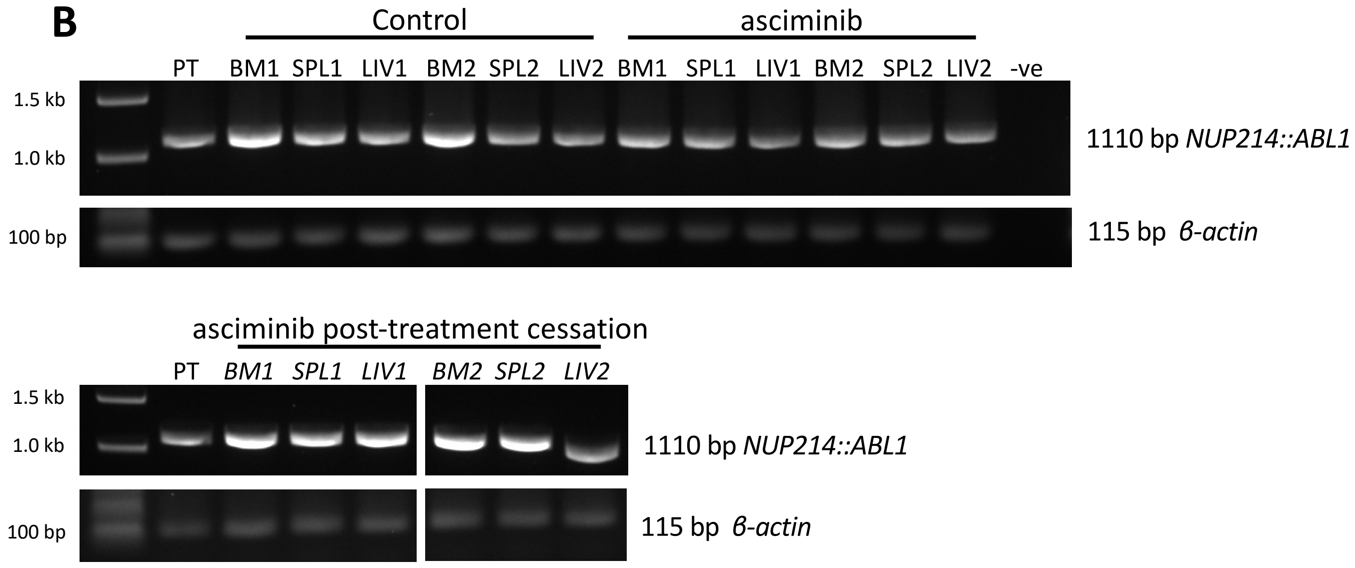

B
Control
asciminib
PT BM1 SPL1 LIV1 BM2 SPL2 LIV2 BM1 SPL1 LIV1 BM2 SPL2 LIV2 -ve
1.5 kb
1.0 kb
1110 bp NUP214::ABL1
100 bp
115 bp β-actin
asciminib post-treatment cessation
PT BM1 SPL1 LIV1 BM2 SPL2 LIV2
1.5 kb
1.0 kb
1110 bp NUP214::ABL1
100 bp
115 bp β-actin


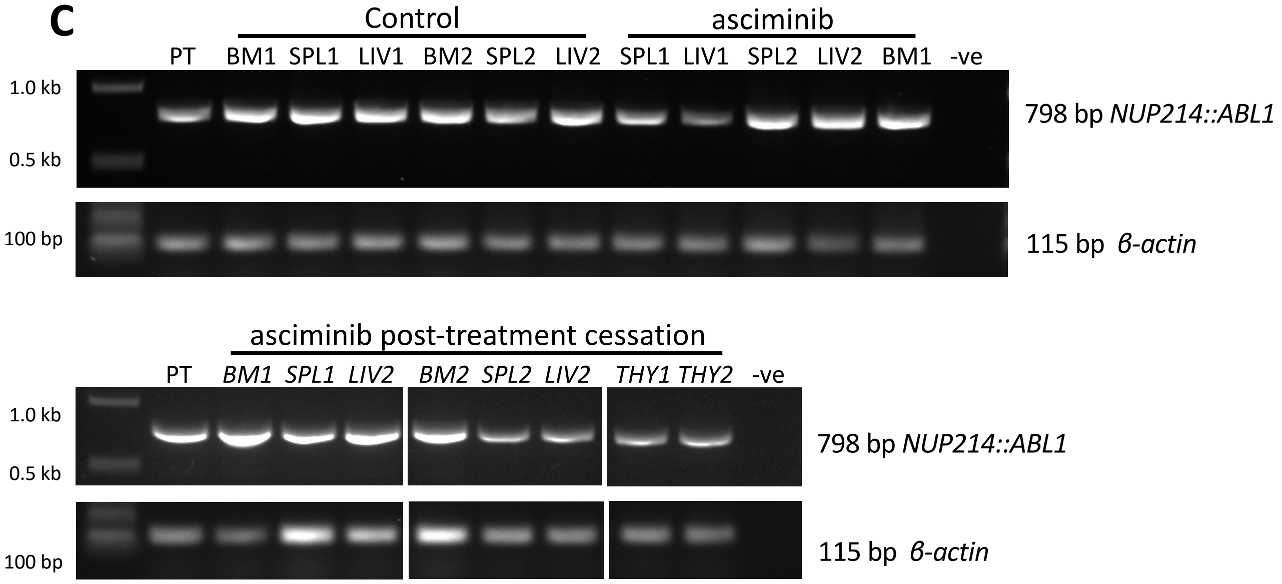

C
Control
asciminib
PT BM1 SPL1 LIV1 BM2 SPL2 LIV2 SPL1 LIV1 SPL2 LIV2 BM1 -ve
1.0 kb
0.5 kb
798 bp NUP214::ABL1
100 bp
115 bp β-actin
asciminib post-treatment cessation
PT BM1 SPL1 LIV2 BM2 SPL2 LIV2 THY1 THY2 -ve
1.0 kb
0.5 kb
798 bp NUP214::ABL1
100 bp
115 bp β-actin

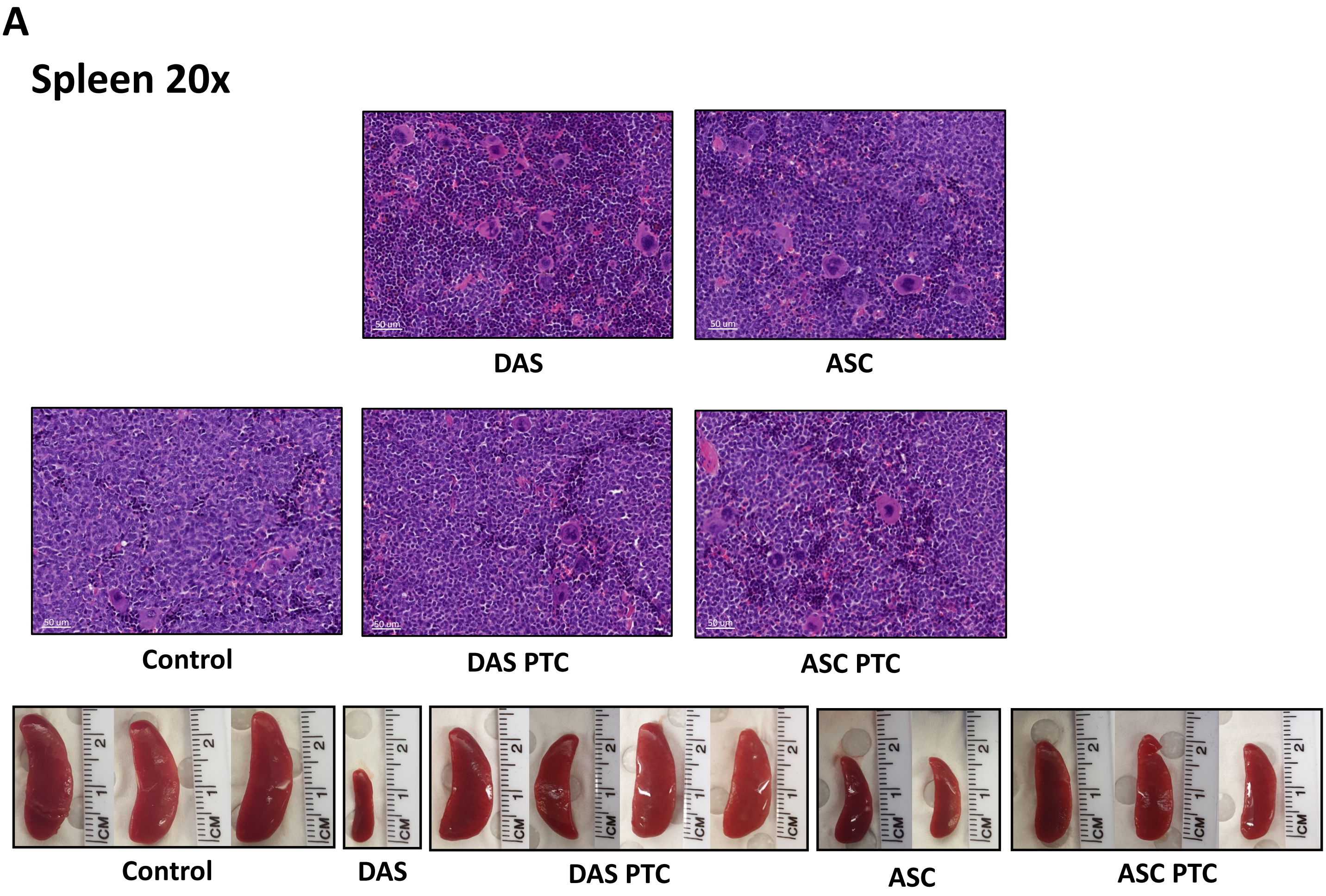


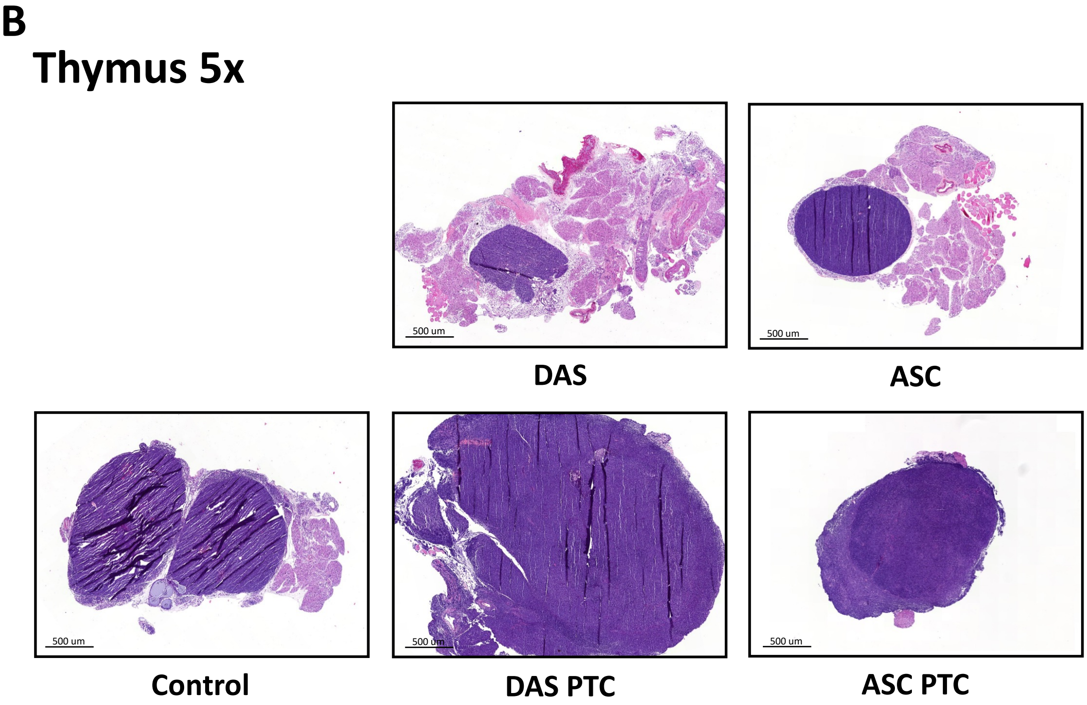


**Supplementary Figure 6. Characterization of leukemic infiltration in a *NUP214::ABL1* PDX model of T-ALL at experimental endpoint, either on treatment or post-treatment cessation.** Organ sections stained with hematoxylin and eosin from *NUP214::ABL1* PDX mice. Images of **A)** spleen **B)** thymus. Scale bars and magnifications are indicated. Pictures of the spleens are included to indicate size differential. Representative images of sections from n=3 mice are shown. DAS=dasatinib, ASC=asciminib.

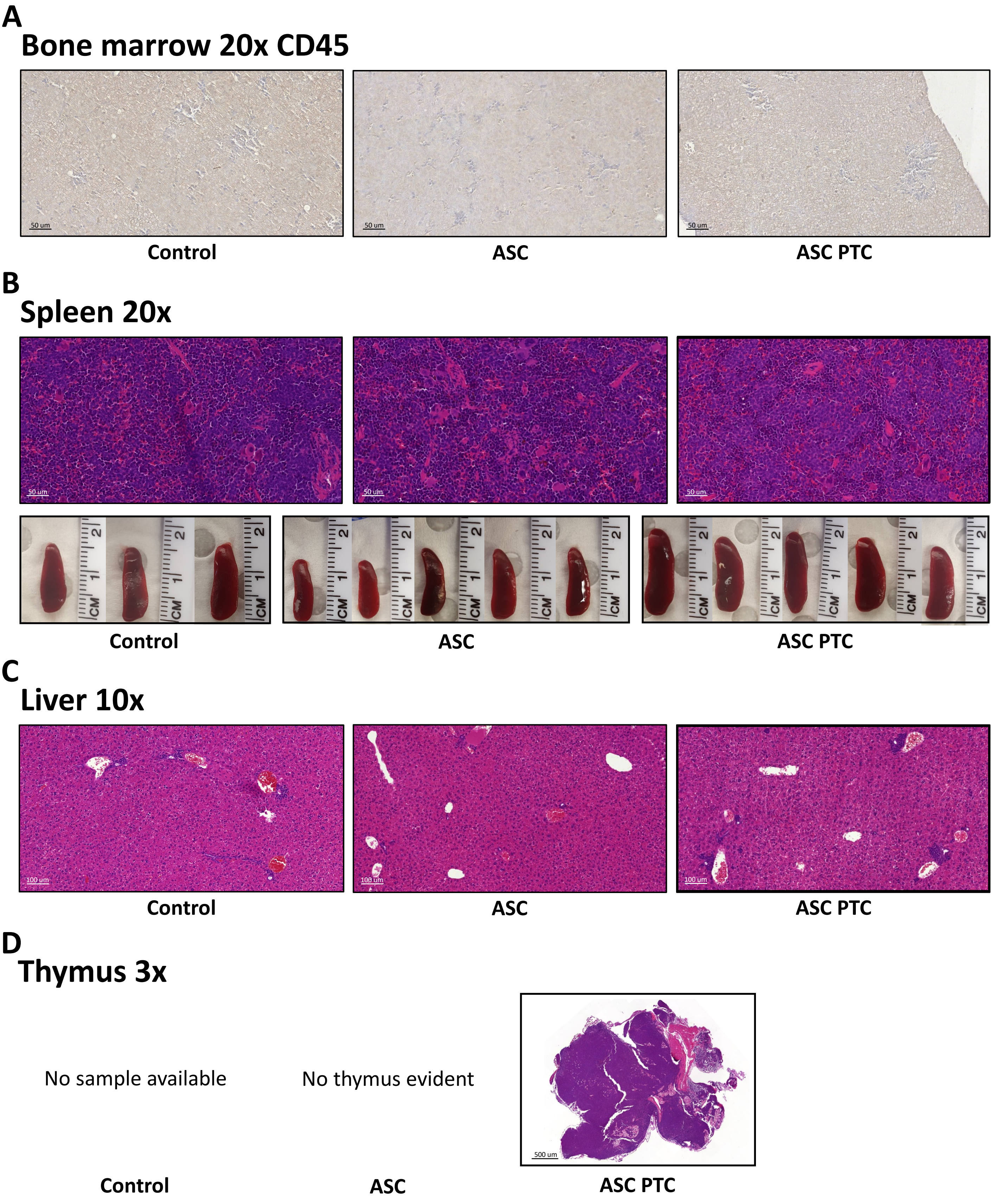


**Supplementary Figure 7. Characterization of leukemic infiltration in a *NUP214::ABL1* PDX model of B-ALL at experimental endpoint, either on treatment or post-treatment cessation.** Organ sections from *NUP214::ABL1* PDX mice engrafted with cells from B-ALL Patient 1. Images from sections stained with hematoxylin and eosin or hCD45 antibody of **A)** bone marrow **B)** spleen **C)** liver **D)** thymus. Scale bars and magnifications are indicated. Pictures of the spleens are included to indicate size differential. Representative images of sections from n=3 mice are shown. ASC=asciminib, PTC=post-treatment cessation.

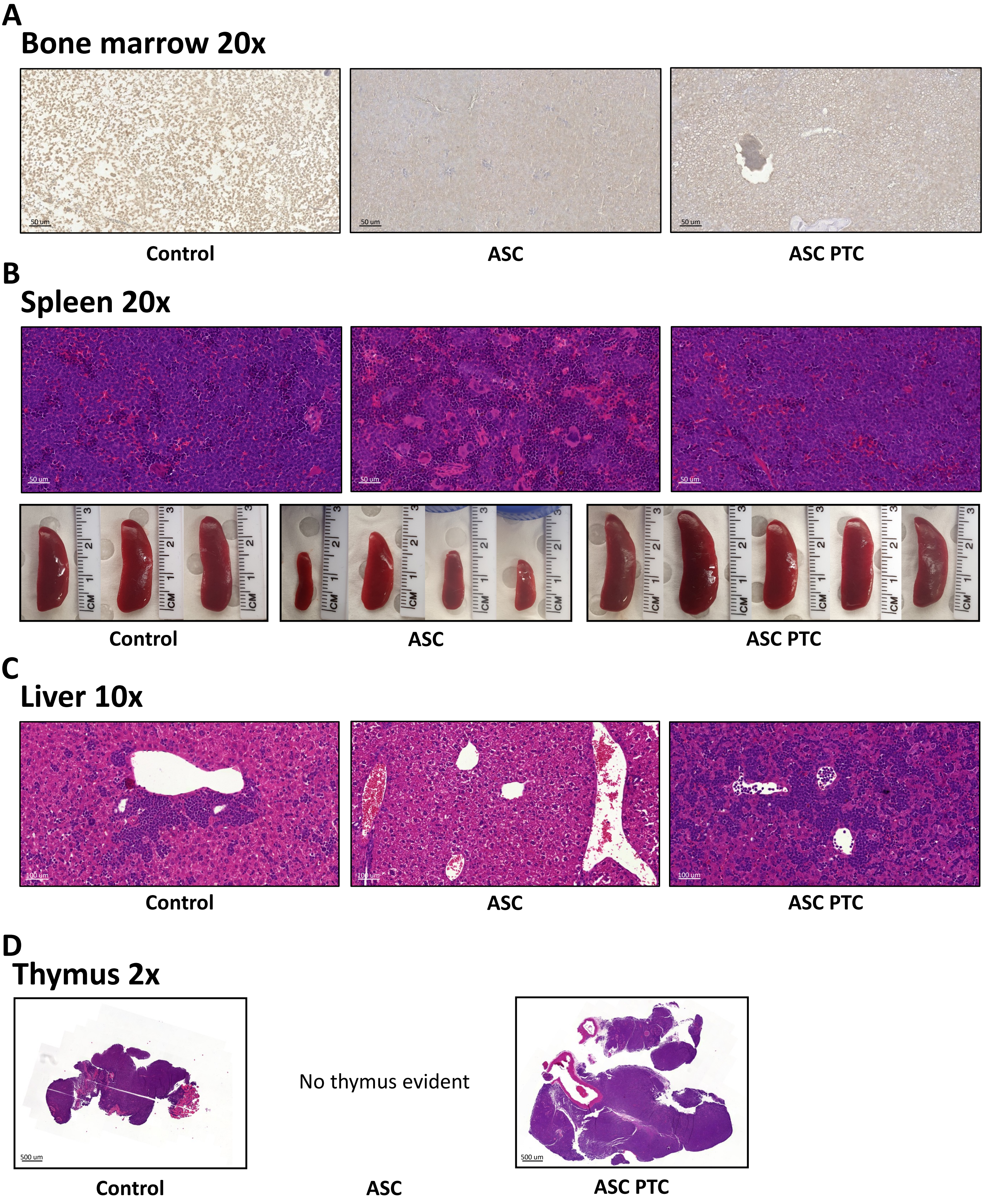


**Supplementary Figure 8. Characterization of leukemic infiltration in *NUP214::ABL1* PDX model of B-ALL at experimental endpoint, either on treatment or post-treatment cessation.** Organ sections from *NUP214::ABL1* PDX mice engrafted with cells from B-ALL Patient 2. Images from sections stained with hematoxylin and eosin or hCD45 antibody of **A)** bone marrow **B)** spleen **C)** liver **D)** thymus. Scale bars and magnifications are indicated. Pictures of the spleens are included to indicate size differential. Representative images of sections from n=3 mice are shown. ASC=asciminib, PTC=post-treatment cessation.

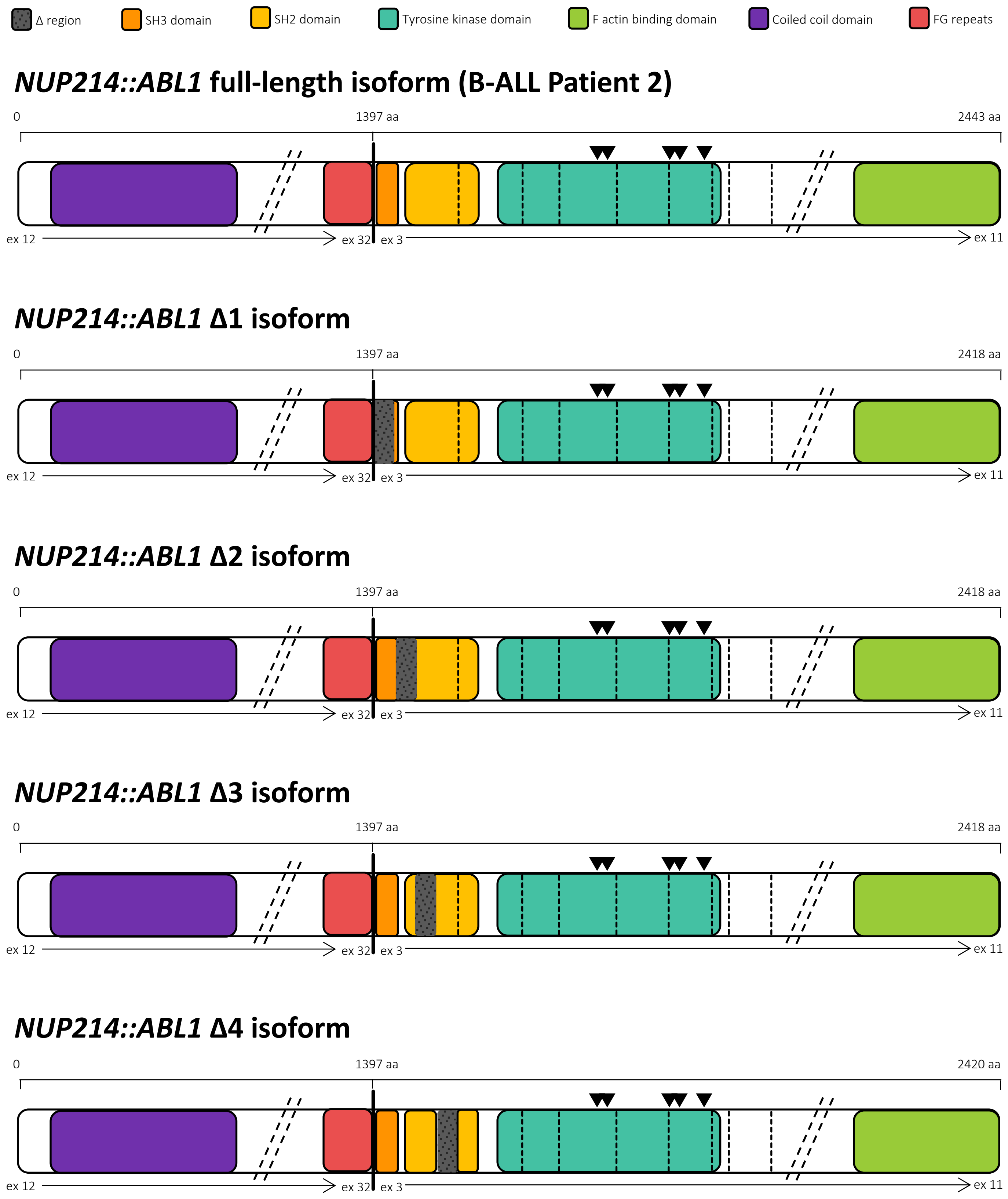


**Supplementary Figure 9. *NUP214::ABL1* fusion gene isoforms depicting the functional domains and exon 3 deletions.** In all isoforms, the residues required for formation of the myristate binding pocket are retained indicated by black triangles above the kinase domain (▼). Deletion of sequential ~25 amino acid regions from within *ABL1* exon 3 are denoted by grey boxes. Breakpoints are denoted by vertical bold black lines and exons by vertical dashed lines. Diagonal dashed lines indicate unstructured regions of genes that have been omitted for simplicity. The *NUP214* NM_005085 and *ABL1* NM_007313 isoforms were used for schematic construction.

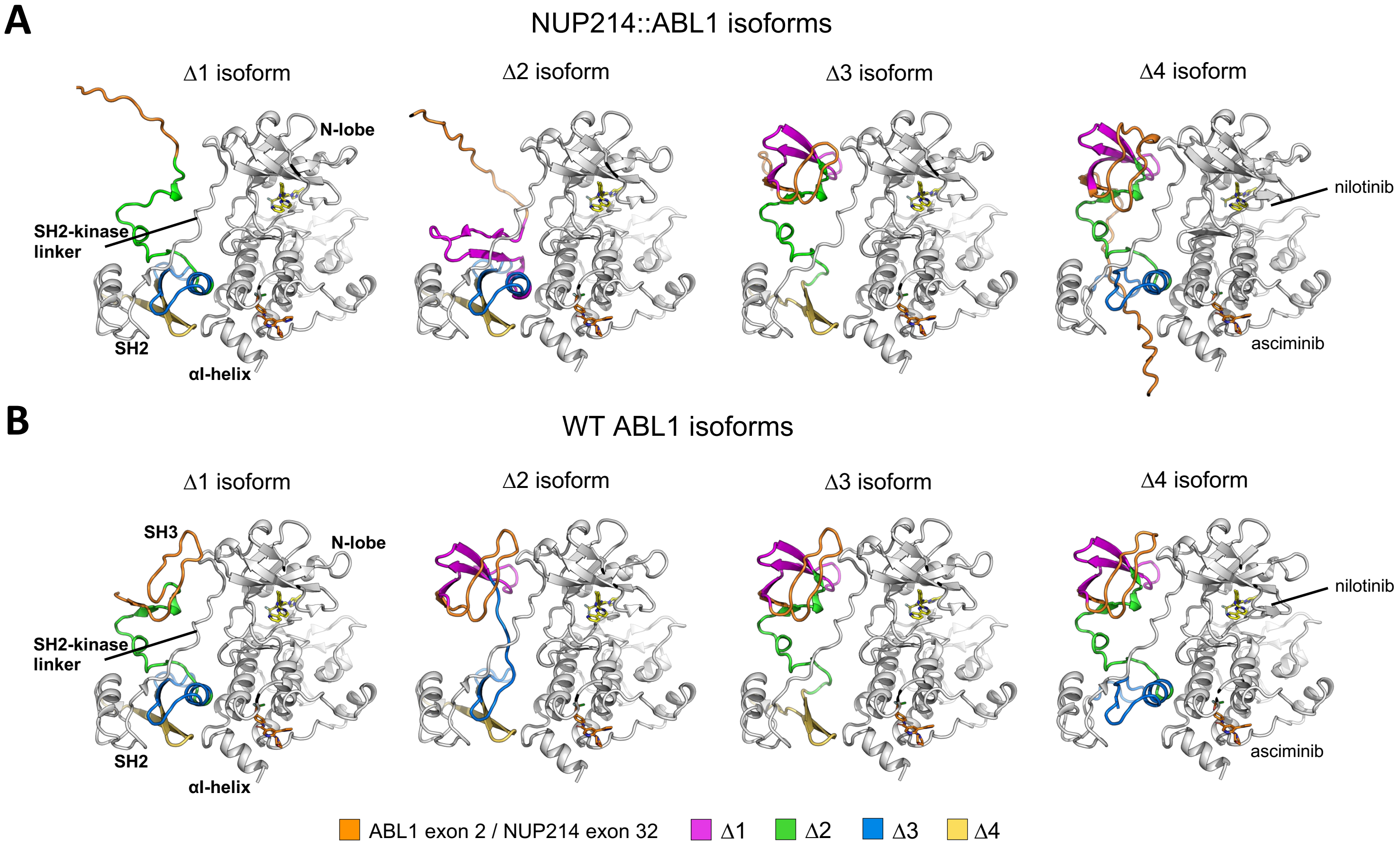


**Supplementary Figure 10. Comparison of predicted structural models of Δ1-4 isoforms within NUP214::ABL1 (e32a3) versus wildtype ABL1 (a2-a10). A)** NUP214::ABL1 (e32a2) Δ1-4 isoforms (as in Figure 6) and **B)** equivalent perspective for wildtype ABL1 highlighting the effects of each deletion region on the interactions between the SH3 domain, the SH2 domain, the kinase domain, and protein folding. The domains and key features are detailed on the Δ1 isoform only and asciminib (orange) and nilotinib (yellow) are shown binding to the myristate and ATP-binding pockets, respectively.

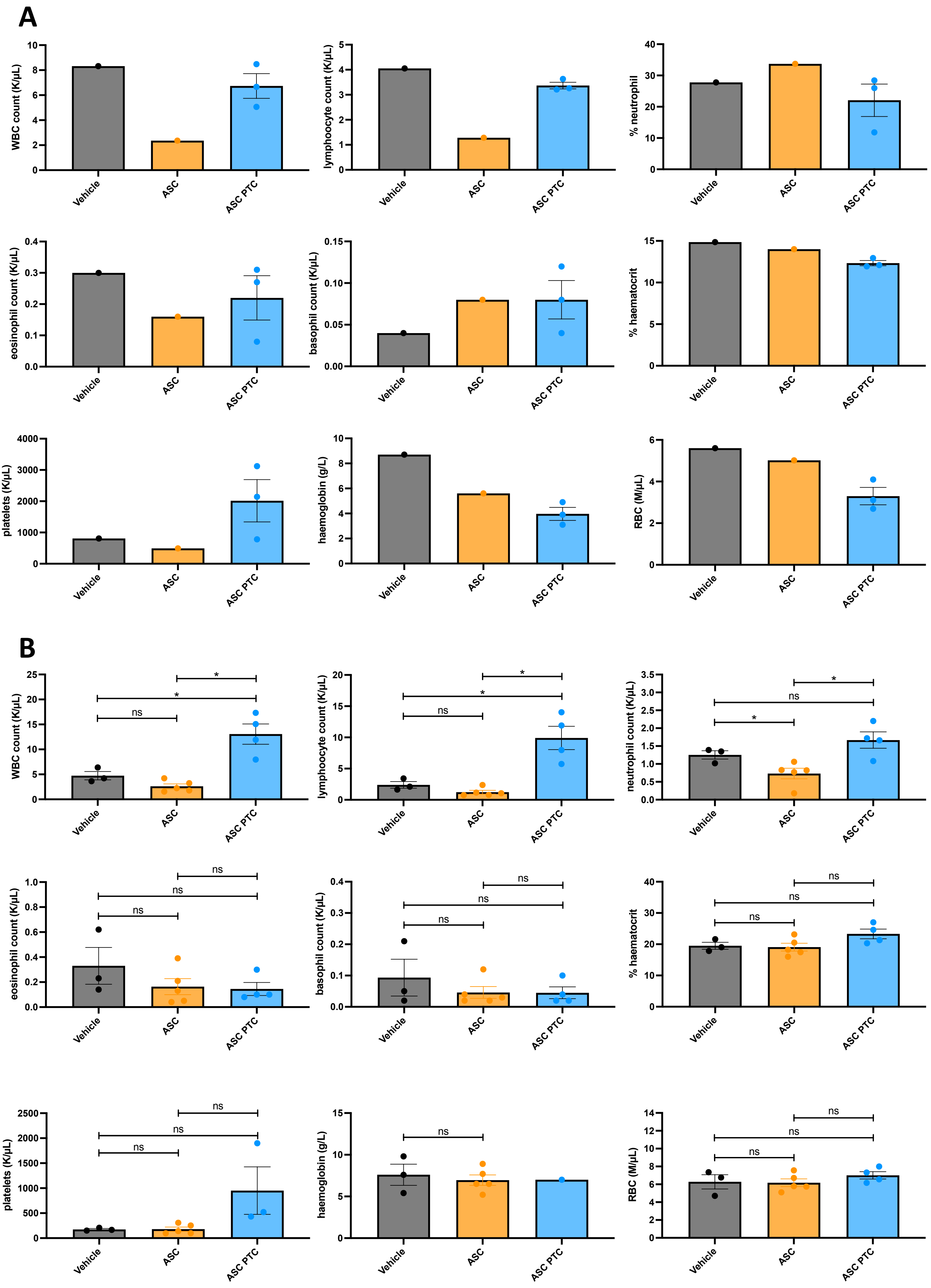
A
WBC count (K/μL)
lymphoocyte count (K/μL)
% neutrophil
eosinophil count (K/μL)
basophil count (K/μL)
% haematocrit
platelets (K/μL)
haemoglobin (g/L)
RBC (M/μL)
Vehicle
ASC
ASC PTC
B
WBC count (K/μL)
lymphoocyte count (K/μL)
neutrophil count (K/μL)
eosinophil count (K/μL)
basophil count (K/μL)
% haematocrit
platelets (K/μL)
haemoglobin (g/L)
RBC (M/μL)
ns
*
Vehicle
ASC
ASC PTC

C

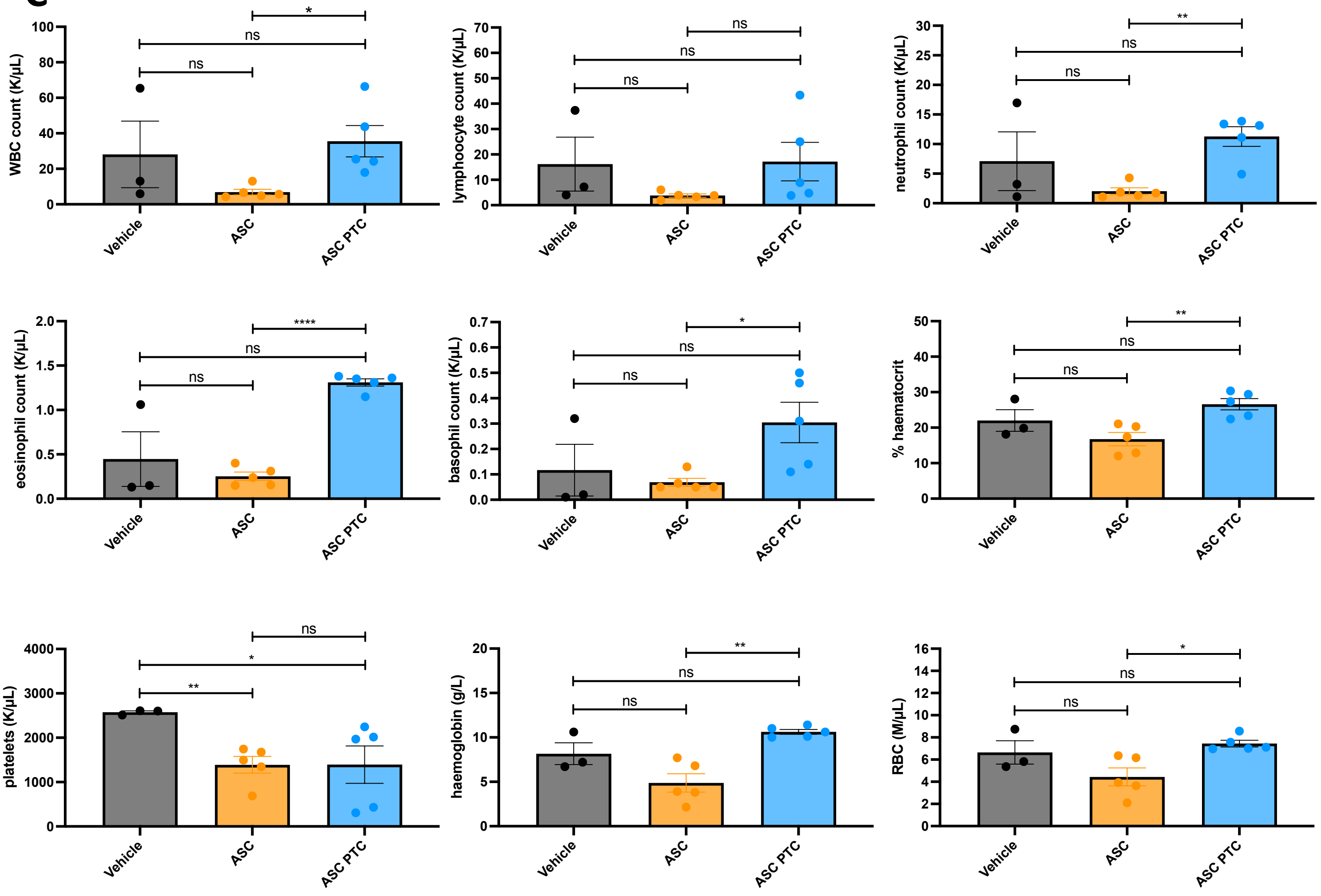


**Supplementary Figure 11. Treatment with asciminib normalizes complete blood counts (CBC) of *NUP214::ABL1* PDX mice at experimental endpoint.** CBC was performed on peripheral blood at experimental endpoint of mice treated with asciminib and CBC cell composition was compared when mice were either on treatment or following treatment cessation. **A)** T-ALL Patient **B)** B-ALL Patient 1 **C**) B-ALL Patient 2. Columns represent mean ±SEM error bars with individual data points shown**.** Welch's unpaired t-test was used to determine significance *p<0.05, **p<0.01, ****p<0.0001. ASC=asciminib, PTC=post-treatment cessation, WBC=white blood cells, RBC=red blood cells.

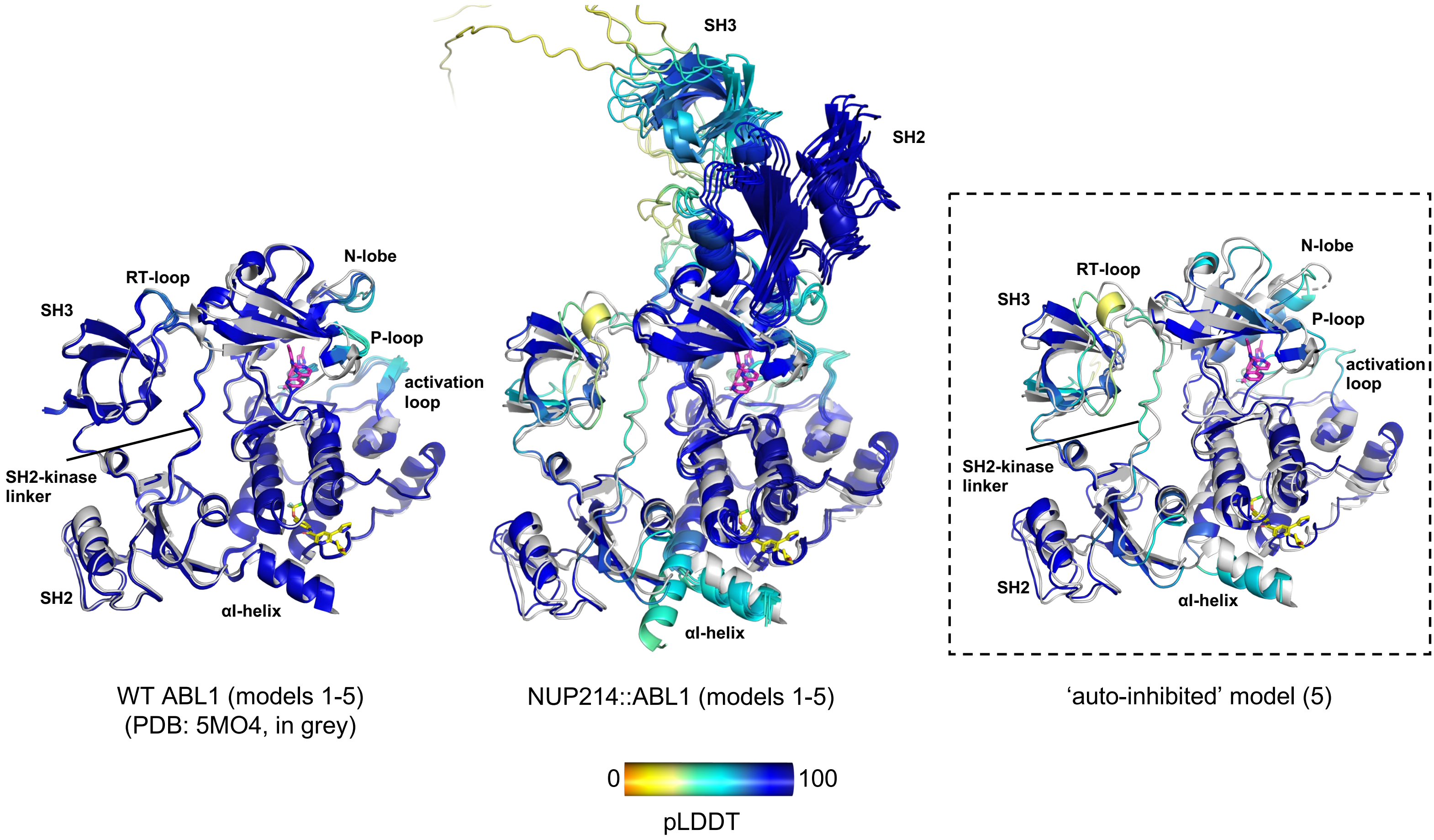


**Supplementary Figure 12. Comparison of the predicted AlphaFold3 models of ABL1 and the NUP214::ABL1 fusion to the X-ray crystal structure of asciminib-bound ABL1.** Wildtype (WT) ABL1 (PDB: 5MO4) in the 'inactive' conformation is shown in grey and all AlphaFold3 models are coloured by predicted local distance difference test (pLDDT) score with higher scores (blue and cyan) indicating higher confidence of the predicted structure. Left: The five AlphaFold3 output models of WT ABL1 aligned to the X-ray crystal structure 5MO4 showing high confidence. Middle: The five AlphaFold3 output models of the NUP214::ABL1 fusion. In four out of the five models, the fusion adopts the 'active' conformation with the SH2 and SH3 domains on top of the kinase-domain N-lobe. Right: The fifth 'auto-inhibited' model of the NUP214::ABL1 fusion aligned to the X-ray crystal structure 5MO4, indicating predicted homology of quaternary structure of NUP214::ABL1 and inactive WT ABL1. Model 5 was selected for further analysis. Asciminib (yellow) and nilotinib (pink) are shown binding to the myristate and ATP-binding pockets, respectively.

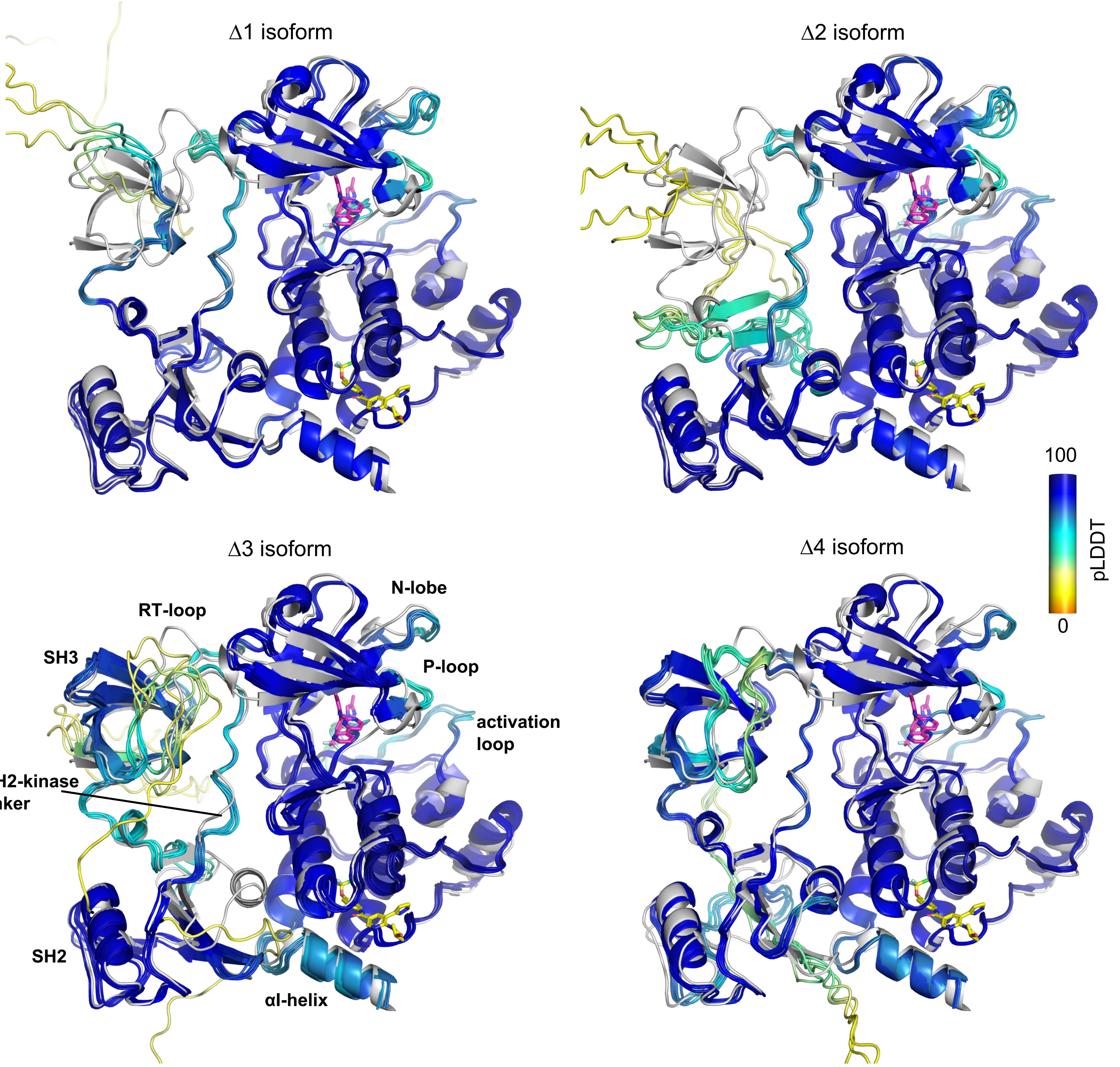


**Supplementary Figure 13. Alignment of the five AlphaFold3 models of NUP214::ABL1 Δ1-4 deletion isoforms to the auto-inhibited ABL1 crystal structure bound to asciminib.** Wildtype (WT) ABL1 (PDB: 5MO4) in the 'inactive' conformation is shown in grey and all AlphaFold3 models are coloured by predicted local distance difference test (pLDDT) score with higher scores (blue and cyan) indicating higher confidence of the predicted structure. Importantly, all five predicted models are in the same architecture for each of the deletion isoforms, with only minor differences between each model. The domains and key features are detailed on the Δ3 isoform only and asciminib (yellow) and nilotinib (pink) are shown binding to the myristate and ATP-binding pockets, respectively.

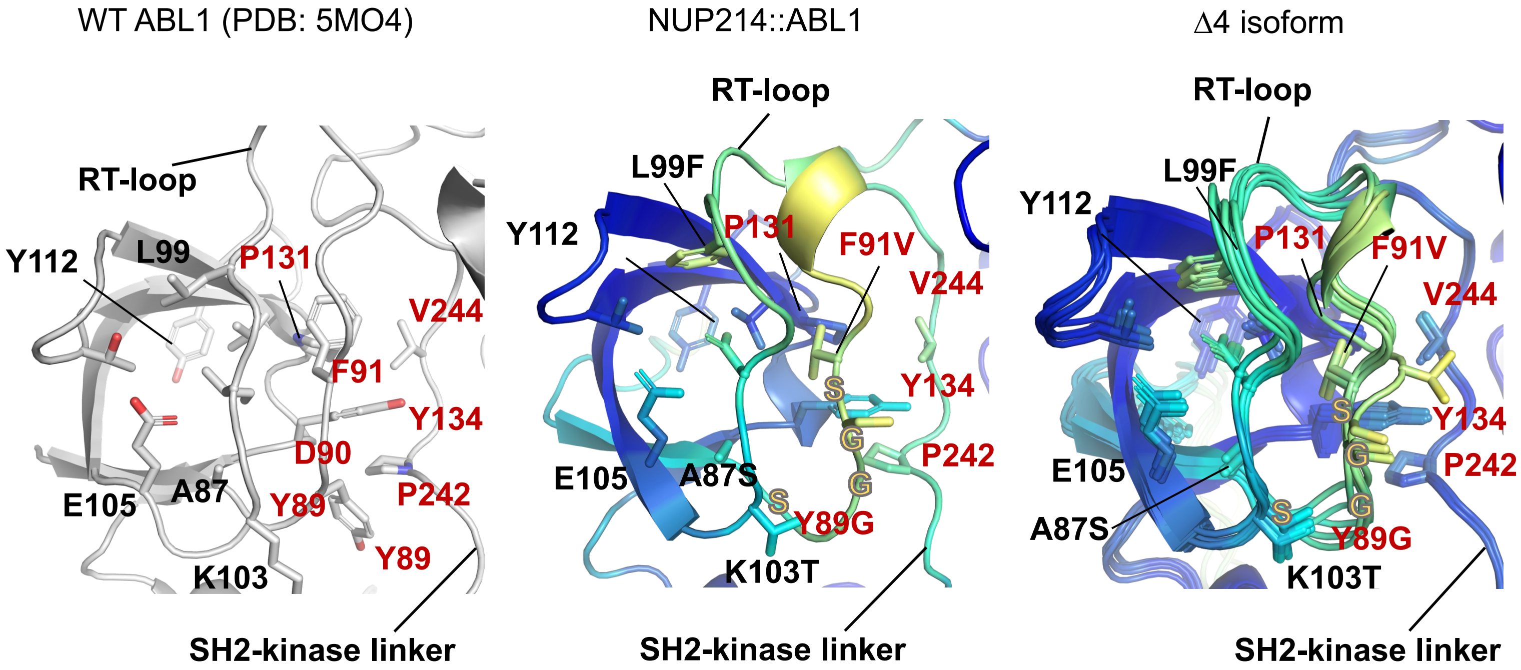


**Supplementary Figure 14. Close-up view of the ABL1 SH3 domain RT-loop in WT ABL1 (left; PDB:5MO4), the full length NUP214::ABL1 fusion (middle) and the Δ4 isoform (right).** In the NUP214::ABL1 fusion and Δ3-4 isoforms, p.Y112 and p.Y134 are located in the same positions and maintain the same interactions with the SH2-kinase linker and RT loop as observed in WT ABL1. However, p.Y89 is replaced by a glycine in NUP214::ABL1 which is also involved in a SSGGGS insertion making the base of the RT-loop more flexible. Residue sidechains are shown as sticks; those comprising the hydrophobic cluster are labelled in red. In the fusion models, residues are annotated by their WT ABL1 numbering and by their substitutions resulting from the fusion as applicable. Additional residues comprising the flexible SSGGGS insertion are shown in orange. AlphaFold3 models are coloured by predicted local distance difference test (pLDDT) score with higher scores (blue and cyan) indicating higher confidence of the predicted structure.
