## Supplementary Notes for "The 5’ fusion partner modulates response to the STAMP inhibitor asciminib in *ABL*-rearranged ALL"

Running title: Asciminib effectively targets *NUP214::ABL1* ALL

Laura N Eadie[1,2], Daniel P McDougal[3,4], Elias Lagonik[1,3], Caitlin E Schutz[1], Tace S Conlin[1,2], Elyse C Page[1,2], Jacqueline Rehn[1,2], John B Bruning[3,4], Andrew S Moore[5,6,7], David T Yeung[1,2,8,9], Timothy P Hughes[1,2,8,9], *Deborah L White[1,2,7,8].

*corresponding author

[1]Precision Cancer Medicine Theme, South Australian Health and Medical Research Institute, Adelaide, Australia

[2]Faculty of Health and Medical Sciences, The University of Adelaide, Adelaide, Australia

[3]Faculty of Sciences, Engineering and Technology, The University of Adelaide, Adelaide, Australia

[4]Institute for Photonics and Advanced Sensing (IPAS), University of Adelaide, Adelaide, Australia

[5]Oncology Service, Children's Health Queensland Hospital & Health Service, Brisbane, Australia

[6]Child Health Research Centre, The University of Queensland, Brisbane, Australia

[7]Australian and New Zealand Children's Haematology-Oncology Group, Victoria, Australia

[8]Australasian Leukemia and Lymphoma Group (ALLG), Melbourne, Australia

[9]Haematology Department, Royal Adelaide Hospital, Adelaide, Australia

**Corresponding Author:**

Professor Deborah White

Theme Director; Precision Cancer Medicine Theme (PCMT).

Group Leader ALL Genomics and Functional Genomics

South Australian Health and Medical Research Institute (SAHMRI)

PO Box 11060, Adelaide South Australia 5001

**Supplementary Note 1 – asciminib treatment of *NUP214::ABL1* mice resolved the leukemic phenotype in the bone marrow**

Immunophenotyping of the primary *NUP214::ABL1* T-ALL patient PB sample revealed 96% CD7+, 58% CD34+ and 40% CD33+ with negligible expression of CD3 and CD13. The vehicle *NUP214::ABL1* PDX mice demonstrated a similar phenotype in the BM with the notable loss of CD34 on the leukemic clone indicating a more mature immunophenotype (Table S3a). Engraftment selected for the most aggressive leukemic clone, with 72±13% hCD7+, 68±15% hCD33+ cells and low/negligible expression of hCD3, hCD13, hCD34. Treatment with dasatinib resulted in the greatest impact on immunophenotype, reducing the hCD7+ population to 6%, the hCD33+ population to negligible expression and increasing the hCD3+ population to 11%. Treatment with asciminib had a similar effect on immunophenotype, reducing hCD7+ and hCD33+ cell populations, while increasing the hCD3+ population. As anticipated, PTC mice receiving asciminib experienced an aggressive outgrowth of the leukemic clone with high hCD7+ and hCD33+ populations (Table S3a). Complete blood examination revealed the total WBC counts were decreased during treatment and increased following treatment cessation (Figure S11A).

Similar results were obtained for both *NUP214::ABL1* B-ALL PDX models. Both *NUP214::ABL1* B-ALL patients initially expressed high levels of CD10/CD19+ cells (Patient 1: 96%, Patient 2: 93%) and CD34+ cells (Patient 1: 95%, Patient 2: 82%). The PDX models demonstrated similar immunophenotype in the vehicle mice (Table S3b). As observed in the T-ALL PDX model, asciminib treatment had a greater effect on the leukemic cells in the PB reducing levels of CD10/CD34+ (Patient 1: 64% to 56±16%; Patient 2: 92±6.6% to 75±14%), CD19/CD34+ (Patient 1: 61% to 14±16%; Patient 2: 88±9.1% to 75±13%) and CD10/CD19+ (Patient 1: 59% to 9.3±13%; Patient 2: 88±8.3% to 72±11%) cells compared with vehicle. PTC, levels of circulating leukemic cells increased to levels comparable to those in vehicle mice. Treatment with asciminib also reduced WBC and lymphocyte counts to within normal ranges (2.6±1.1 K/μL and 1.2±0.7 K/μL, respectively; Fig S11). However, upon asciminib cessation, a significant increase in total WBC (Patient 1: 13±4.1 K/μL, p=0.0117; Patient 2: 35.5±20 K/μL, p=0.0302) and lymphocyte (17±17 K/μL) counts was observed. Haemoglobin, haemocrit and RBC levels were not reduced in mice receiving asciminib compared with vehicle (Fig S11B-C).

**Supplementary Note 2 – phosphorylation of STAT5 is reduced with asciminib treatment even following treatment cessation**

It could be argued that the sustained inhibition of kinase signaling following treatment cessation was due to residual drugs still being bound to ABL1. However, this is unlikely given the half-lives of dasatinib and asciminib and the number of intervening days between treatment cessation and experimental end-point sample collection. The standard dose of dasatinib (100 mg per day) has a half-life of ~3-5 hours in adults[1] and a dose of asciminib (80 mg per day) has a half-life of 5.5 hours in adults[2]. Mice in our PDX models were given 20 mg/kg/day dasatinib and 30 mg/kg/day asciminib, equating to 0.4-0.46 mg and 0.57-0.72 mg daily for dasatinib and asciminib respectively (amount of drug depended on the weight of individual mice). The median number of days from treatment cessation to experimental endpoint was 28 days for asciminib-treated mice and 35.5 days for dasatinib-treated mice so negligible amounts of drugs would have been present in the mice at the time of sample collection.

**Supplementary Note 3 – determination of the critical region of the ABL1 SH3 domain required for asciminib efficacy**

PDX models were generated from *NUP214::ABL1* patients with different *ABL1* breakpoints as indicated in Figure S1: the T-ALL patient had an *ABL1* exon 2 breakpoint (complete SH3 domain present), while the two B-ALL patients had *ABL1* exon 3 breakpoints (partial SH3 domain present). Despite the different breakpoints, asciminib demonstrated efficacy in all three PDX models. Additionally, previous *in vitro* work indicated *ABL1* exon 2 was not necessary for asciminib efficacy in an *in vitro NUP214::ABL1* model[3]. So for our current *in vitro* work dissecting the regions of the SH3 essential for asciminib efficacy, we amplified material from B-ALL patient 2 as the *ABL1* exon 3 breakpoint already omitted part of the SH3 domain. The remaining portion of *ABL1* exon 3 (98 amino acids) was then divided into four sections that were individually deleted by site directed mutagenesis (Δ1-3=25 amino acids; Δ4=23 amino acids). In this way, we were able to interrogate the key region of the ABL1 SH3 domain required for asciminib efficacy.

**Supplementary Note 4 – structural modeling of the quaternary architecture of the NUP214::ABL1 fusion and the Δ1-4 isoforms**

Structural models of the NUP214 fusion and Δ1-4 isoforms were predicted using AlphaFold3[4]. Asciminib and TKIs were subsequently modelled using ICM-Pro (Molsoft L.L.C.). The five models generated from AlphaFold3 for wildtype (WT) ABL1, full length NUP214::ABL1 and each of the deletion isoforms were superimposed and coloured according to the predicted local distance difference test (pLDDT) values. Regions of the structures with high pLDDT scores (blue) indicate that the predicted structure has a high confidence of agreeing with the experimental structure while regions with lower pLDDT scores (cyan, yellow) are of lower confidence.

The WT ABL1 model was generally high confidence (>70 pLDDT) and aligned well with the crystal structure of the auto-inhibited state (PDB ID: 5MO4). The regions coloured lower confidence correspond with the activation loop, P-loop, αC-helix and SH2-kinase linker which are known to be conformationally dynamic or flexible in Abl-family kinases[5,6] (Figure S12). The N-terminal 5' region preceding the SH3 domain (RT-loop) is also slightly lower confidence than the surrounding areas reflecting the unstructured/disordered nature of this region.

Among the five predicted models of the full-length NUP214::ABL1, four adopted an active conformation with the SH2 and SH3 domains stacked on the N-lobe of the kinase domain, consistent with the active state architecture[7]. Only one model adopted the auto-inhibited inactive conformation. Given that the inactive conformation occurs upon auto-inhibition by the myristate moiety (or upon drug binding), and the myristate modification cannot be added into the model, this model was selected for further analysis as it best represented conformational state following asciminib inhibition. As in WT ABL1, the activation loop, P-loop, αC-helix, and SH2-kinase linker were predicted with lower confidence. These patterns were broadly consistent across the Δ1-4 isoform models (Figure S13).

In the full-length NUP214::ABL1 fusion and Δ4 isoform, the RT-loop of the SH3 domain adopted a conformation similar to that of WT ABL1, despite being encoded by exon 32 of NUP214, though with markedly lower confidence (Figure S13). In the Δ3 isoform, four out of the five models retained an 'intact' RT-loop, albeit with greater conformational variability. The Δ1 and Δ2 isoforms did not form a folded SH3 domain. This was as expected since the Δ1 isoform lacks an integral part of the SH3 domain with the amino acid sequence GYNHN that contacts the αC-helix on the kinase domain N-lobe (p.G111-N115 as per NM_007313, or p.G92-NN96 as per NM_005157[8] and the Δ2 isoform lacks the SH2-SH3 domain linker (amino acid sequence ITPVN[8] corresponding to p.I135-N139 or p.I116-N120, NM_007313 and NM_005157 isoforms respectively). Instead, the Δ2 isoform was predicted to form two β-sheets stacked on the SH2 domain, albeit with low confidence and/or lack of secondary structure. Residues in these β-sheets are involved in the *de novo* hydrogen bond network discussed in more detail in the main text.

Closer analysis of the RT-loop and SH3 domain in the full-length NUP214::ABL1 fusion and Δ3-4 models revealed that the hydrophobic cluster important for SH3 domain docking to the SH2-kinase linker was preserved, centred around p.Y134 in WT ABL1 (NM_007313, or p.Y115 for NM_005157) (Figure S14)[5,9]. In the fusion, p.Y89 located at the N-terminal of the RT-loop is replaced by a flexible SSGGGS insertion. Despite this substitution, backbone and side-chain interactions suggest potential structural retention of the RT-loop and SH3 domain capping. We hypothesize that partial preservation of the SH3 domain in the full-length NUP214::ABL1 fusion and Δ3-4 isoforms may enable SH2-kinase linker docking and support allosteric inhibition of kinase activity by asciminib. However, whether the RT-loop remains structured in these contexts remains unclear and warrants future investigation.

In the full-length NUP214::ABL1 fusion and Δ1-2 isoforms, the SH2 domain was intact and is predicted with high confidence, similar to WT ABL1 model. Notably, despite deletions in the SH2 domain, the Δ3-4 isoform models retained proximity between the truncated SH2 domain and the allosteric αI-helix of the myristate binding pocket. In the Δ4 isoform, exon 32 of the NUP214 fusion extends below the β-strand and is sandwiched between two α-helices, though this region is of notably lower confidence (Figure S13). Nonetheless, our *in vitro* data demonstrate that these specific deletions to the SH2 domain do not impair asciminib sensitivity, suggesting previously uncharacterized plasticity in allosteric regulation of ABL1. Like the RT-loop, the structural implications of these predictions require experimental validation.

**Supplementary References**


1 Dasatinib (Sprycel ®) [Prescribing Information]. In: *Princeton, New Jersey: Bristol Myers Squibb Company* (Revised 07/2024).

2 Asciminib (SCEMBLIX ®) [Prescribing Information]. In: *East Hanover, New Jersey: Novartis Pharmaceuticals Corporation* (Revised 08/2024).

3 Eadie, L. N., Lagonik, E., Page, E. C. *et al.* Asciminib is a novel inhibitor of ABL1 and ABL2 gene fusions in ALL but requires the ABL SH3 domain for efficacy. *Blood* **144**, 1022-1026 (2024).

4 Abramson, J., Adler, J., Dunger, J. *et al.* Accurate structure prediction of biomolecular interactions with AlphaFold 3. *Nature* **630**, 493-500 (2024).

5 Panjarian, S., Iacob, R. E., Chen, S., Engen, J. R. & Smithgall, T. E. Structure and dynamic regulation of Abl kinases. *J Biol Chem* **288**, 5443-5450 (2013).

6 Xie, T., Saleh, T., Rossi, P. & Kalodimos, C. G. Conformational states dynamically populated by a kinase determine its function. *Science* **370** (2020).

7 Nagar, B., Hantschel, O., Young, M. A. *et al.* Structural basis for the autoinhibition of c-Abl tyrosine kinase. *Cell* **112**, 859-871 (2003).

8 Leyte-Vidal, A., DeFilippis, R., Outhwaite, I. R. *et al.* Absence of ABL1 exon 2-encoded SH3 residues in BCR::ABL1 destabilizes the autoinhibited kinase conformation and confers resistance to asciminib. *Leukemia* 10.1038/s41375-024-02353-0 (2024).

9 Manley, P. W., Barys, L. & Cowan-Jacob, S. W. The specificity of asciminib, a potential treatment for chronic myeloid leukemia, as a myristate-pocket binding ABL inhibitor and analysis of its interactions with mutant forms of BCR-ABL1 kinase. *Leukemia research* **98**, 106458 (2020).
