## Supplementary Table 1 for "The 5’ fusion partner modulates response to the STAMP inhibitor asciminib in *ABL*-rearranged ALL"

| Sample ID | Disease | Gender | Sample | sampleType | Age at Dx | Age at REL | Fusion | Breakpoint of most abundant fusion isoform |
|---|---|---|---|---|---|---|---|---|
| ADI_0393 | T-ALL | F | REL2 | PB | not known | 54 | *NUP214::ABL1* | *NUP214(exon 31)::ABL1(exon2)* |
| CHI_0235 | B-ALL Patient 1 | M | Dx | BM | 15 | relapse status unknown | *NUP214::ABL1* | *NUP214(exon 34)::ABL1(exon3)* |
| QCTB0709_1617303 | B-ALL Patient 2 | M | Dx | BM | 12 | relapse status unknown | *NUP214::ABL1* | *NUP214(exon 32)::ABL1(exon3)* |

NOTE: A DNA sample was not available for B-ALL Patient 1 so MLPA has been performed on total white cells from a PDX BM sample instead (96.1% hCD45 and 95.3% hCD19)

Abbreviations: Dx=diagnosis; REL=relapse; PB=peripheral blood; BM=bone marrow; WBC=white blood cell; MLPA=Multiplex ligation-dependent probe amplification; PDX=patient-derived xenograft

| Sample ID | Disease | Cytogenetics | Additional key lesions: MLPA | Additional key lesions: mRNA-Seq |
|---|---|---|---|---|
| ADI_0393 | T-ALL | Two leukemic clones were identified.<br>46,XX,t(4;9)(q?32-34;q?31-33) [12]<br>46,s1,del(17)(p11.2) [3]<br>46,XX [16] | *IKZF1* heterozygous deletion (exons 1-8)<br>*CDKN2A* homozygous deletion (exons 2 and 4, entire gene)<br>*CDKN2B* homozygous deletion (exons 1 and 2, entire gene)<br>*MTAP* heterozygous deletion (exon 1)<br>*ABL1* duplication (exons 4-12)<br>*NUP214* duplication (exons 2-23)<br>*PHF6* homozygous duplication (exons 1-10, entire gene) | no additional lesions of clinical relevance |
| CHI_0235 | B-ALL Patient 1 | 46XY [30] | *IKZF1* heterozygous deletion (intron 3, exons 5 and 7)<br>*IKZF1* sub-clonal deletion (exons 4 and 6)<br>No CNVs in *IKZF1* exons 1-3 and 8<br>*IGHD-up* homozygous deletion<br>*ETV6* heterozygous deletion (exons 2-8, not exon 1)<br>*ERG* sub-clonal duplication (intron 3, exons 5 and 8, not exons 1-3,4,6-7,9-12)<br>*PAX5* sub-clonal duplication (exons 1-2 and 6-8, not exons 5 or 10)<br>*ABL1* duplication (exons 4-12)<br>*NUP214* duplication (exons 2-23)<br>*PTEN* sub-clonal duplication (exons 1 and 9) | *EZH2* p.D608G/p.D664G (VAF=0.33) |
| QCTB0709_1617303 | B-ALL Patient 2 | Two unrelated leukemic clones were identified.<br>46,XY,del(4)(p11),add(15)(q22),add(19)(q13)[8]<br>45,XY,add(18)(p11),-20[8]<br>46,XY[2]<br>nuc ish(PBX1x2,TCF3x1)[62/200]<br>PTPRT (20q12) and MYBL2(20q13.12)x1[76/200] | *IKZF1* heterozygous deletion (exons 4-7, not exons 1-3)<br>*EBF1* heterozygous deletion (exons 1-16, entire gene)<br>*PAX5* heterozygous deletion (exons 1-6, not exons 7-10)<br>*BTG1* heterozygous deletion (exon 2, not exon 1)<br>*ABL1* duplication (exons 4-12)<br>*NUP214* duplication (exons 2-23)<br>*PTPN2* subclonal deletion (exons 1-9, entire gene) | no additional lesions of clinical relevance |
