## Supplementary Table 2 for "The 5’ fusion partner modulates response to the STAMP inhibitor asciminib in *ABL*-rearranged ALL"

Table S2a: Base pair and primer sequences for *NUP214::ABL1* SDM1-4 isoforms

| | Base pair sequence as per NM_007313 | FORWARD primer (5' → 3') | REVERSE primer (5' → 3') |
|---|---|---|---|
| SDM1 | GAAAAGCTCCGGGTCTTAGGCTATAATCACAATGGGGAATGGTGTGAAGCCCAAACCAAAAATGGCCAAGGCTGG | GTCCCAAGCAACTACATC | ACCTGTTTTGTTTGGAGAG |
| SDM2 | GTCCCAAGCAACTACATCACGCCAGTCAACAGTCTGGAGAAACACTCCTGGTACCATGGGCCTGTGTCCCGCAAT | GCCGCTGAGTATCTGCTG | CCAGCCTTGGCCATTTTTG |
| SDM3 | GCCGCTGAGTATCTGCTGAGCAGCGGGATCAATGGCAGCTTCTTGGTGCGTGAGAGTGAGAGCAGTCCTGGCCAG | AGGTCCATCTCGCTGAGATACG | ATTGCGGGACACAGGCCC |
| SDM4 | AGGTCCATCTCGCTGAGATACGAAGGGAGGGTGTACCATTACAGGATCAACACTGCTTCTGATGGCAAG | CTCTACGTCTCCTCCGAG | CTGGCCAGGACTGCTCTC |

Table S2b: Optimsed cycling conditions for site-directed mutagenesis

| Cycling conditions | SDM1 | SDM2 | SDM3 | SDM4 |
|---|---|---|---|---|
| Initial denaturation | 98$^{o}$C, 30 seconds | | | |
| 25 cycles | 98$^{o}$C, 10 seconds | | | |
| | 63$^{o}$C, 30 seconds | 69$^{o}$C, 30 seconds | 72$^{o}$C, 30 seconds | 68$^{o}$C, 30 seconds |
| | 72$^{o}$C, 8 minutes | | | |
| Final extension | 72$^{o}$C, 2 minutes | | | |
| Hold | 4$^{o}$C, ∞ | | | |
