## Supplementary Table 3 for "The 5’ fusion partner modulates response to the STAMP inhibitor asciminib in *ABL*-rearranged ALL"

**Table S3a. Immunophenotyping of bone marrow from mice in a *NUP214::ABL1* PDX model of T-ALL, either on treatment or post treatment cessation.**

| | hCD7+/hCD34+ | hCD33+/hCD34+ | hCD3+/hCD34+ | hCD7+/hCD3+ |
|---|---|---|---|---|
| **Patient PBMNCs** | 53 | 21 | 1.3 | 2.6 |
| | **hCD7+/hCD34-** | **hCD33+/hCD34-** | **hCD3+/hCD34-** | **hCD7+/hCD3+** |
| **Vehicle control** | 72±13 | 68±15 | 2.0±1.5 | 0.7±0.4 |
| **DAS** | 6.3 | 0 | 11 | 0 |
| **ASC** | 60±0.6 | 44±16 | 9.6±1.1 | 4.1±0.9 |
| **ASC PTC** | 87 | 88 | 4.7 | 5.8 |

Values represent cell percentage of total population of live cells analyzed via flow cytometry. PBMNCs=peripheral blood mononuclear cells, DAS=dasatinib, ASC=asciminib, PTC=post treatment cessation.

**Table S3b. Immunophenotyping of peripheral blood from mice in a *NUP214::ABL1* PDX models of B-ALL, either on treatment or post treatment cessation.**

| | hCD10+/hCD19+ | hCD10+/hCD34+ | hCD19+/hCD34+ |
|---|---|---|---|
| **Patient 1 PBMNCs** | 96 | 94 | 94 |
| **Vehicle control** | 59 | 64 | 61 |
| **ASC** | 9.3±13 | 56±16 | 14±16 |
| **ASC PTC** | 62±28 | 70±27 | 63±26 |
| **Patient 2 PBMNCs** | 93 | ND | 82 |
| **Vehicle control** | 88±8.3 | 92±6.6 | 88±9.1 |
| **ASC** | 72±11 | 75±14 | 75±13 |
| **ASC PTC** | 88±7.8 | 89±8.8 | 88±7.9 |

Values represent cell percentage of total population of live cells analyzed via flow cytometry. PBMNCs=peripheral blood mononuclear cells, ASC=asciminib, PTC=post treatment cessation, ND=not determined.
